# Engineering Cell Sensing and Responses Using a GPCR-Coupled CRISPR-Cas System

**DOI:** 10.1101/152496

**Authors:** P. C. Dave P. Dingal, Nathan H. Kipniss, Louai Labanieh, Yuchen Gao, Lei S. Qi

## Abstract

G-protein coupled receptors (GPCRs) are the largest and most diverse group of membrane receptors in eukaryotes, and detects a wide array of physiological cues in the human body. We describe a new molecular device that couples CRISPR-Cas9 programmed genome regulation to natural and synthetic extracellular signals via GPCRs. The design of our synthetic device, named CRISPR ChaCha, displays superior performance over an architecture proposed by the previously reported Tango system. Using a parsimonious mathematical model and gene-reporter assays, we find that CRISPR ChaCha can recruit and activate multiple Cas9 molecules for each GPCR molecule. We also characterize key molecular features that modulate CRISPR ChaCha performance. We adopt the design to diverse GPCRs that sense synthetic and natural ligands including chemokines, mitogens, and fatty acids, and observe efficient conversion of signals to customizable genetic programs in mammalian cells, including regulation of endogenous genes. The new class of CRISPR-coupled GPCRs provides a robust and efficient platform for engineering cells with novel behaviors in response to the diverse GPCR ligand repertoire.

---

Eukaryotic cells have evolved diverse classes of transmembrane receptors to transduce various extracellular signals into intracellular responses. Binding of receptors and their cognate ligands triggers their intracellular enzymatic activities. Inside the cell, this leads to complex signaling of downstream signaling protein and secondary messenger functions that ultimately transduce signals to genomic programs. The complexity of these signaling cascades has made it a major challenge to harness natural signaling pathways for cell engineering in a flexible and programmable manner [1].

In the interest of designing rational control over how cells generate behaviors in response to environmental cues, work has been done in cell receptor engineering. Customized cell sensing/response pathways based on synthetic receptors, such as the chimeric antigen receptor (CAR) [2] and the synthetic Notch (synNotch) receptor [3], have been used to engineer T cells to respond to certain disease-relevant signals [4]. These receptors employ antibody-based extracellular domains (e.g., single-chain antibodies or nanobodies), which are limited by the availability of such domains and can only sense cell-surface proteins. Other works developed self-contained receptor–signal transduction systems [5-7].

An important concept in cell signaling biology is signal amplification, where the many downstream molecules can be activated or released for one receptor-ligand complex [8]. Much of the previous work has designed “one molecule in, one molecule out” systems, and do not lead to significant signal amplification thereby limiting utility [3, 5-7]. Beyond these systems, engineering modular and efficient synthetic receptors for mammalian cells would benefit from a generalizable engineering approach that can not only expand the ligand repertoire (e.g., to soluble peptides, chemokines, fatty acids, among many others) but also convert these cues into customizable transcriptional programs.

We wanted to activate downstream genes in a manner than is amenable to different receptor inputs. G-coupled protein receptors (GPCRs) are attractive candidates for receptor engineering because they can detect diverse classes ligands including endogenous hormones and growth factors, and natural or synthetic small molecules [9-12]. Previous work in receptor engineering replaced the intracellular domains with a proteolytically cleavable artificial transcription factor (e.g., Gal4, rtTA) to create a genetic reporter of receptor activity [3, 6, 7]. However, the efficiency of these systems is limited by the inability to amplify signals or regulate endogenous genes. We thus needed to create a design that could permit signal amplification, but also flexibly target the genome.

CRISPR-Cas9 technologies have revolutionized programmable genome manipulations. Cas9 can be precisely targeted to a genomic locus of interest for gene editing using a customized single guide RNA (sgRNA) [13-15]. Beyond editing, the nuclease-deactivated Cas9 (dCas9) molecule can be fused with effector protein domains to activate or repress transcription of genes of interest or to modify the epigenome [16-21]. In addition, multiple sgRNAs can be used simultaneously regulate different targets and drive complex gene expression programs [18]. The flexibility of CRISPR-Cas systems for genome regulation makes them powerful tools for programming novel sensor-coupled cell behaviors.

Here we explore strategies to repurpose the GPCR receptor as an efficient platform for converting the input-sensing ability of GPCRs into programmed genome responses via CRISPR-dCas9 technologies. Starting with an engineered synthetic GPCR that recognizes a bioinert ligand, we explore two design architectures to search for better ligand detection and gene regulation. We demonstrate that our new design, named CRISPR ChaCha, significantly outperformed the previously reported Tango system [6, 7]. We formulate rate models to explain key design parameters of the CRISPR ChaCha that confer better performance. We adopt this new architecture for additional GPCRs that detect natural and synthetic ligands. We also demonstrate that our system could be used to detect synthetic or natural ligands to activate endogenous gene programs.

## RESULTS

### Implementation of two alternative CRISPR-coupled GPCR designs

In search of more efficient sensor-effector architectures than previously reported, we created two designs for proteolytically coupling dCas9 function to GPCRs (**Fig. 1a**). The first design is derived from a previously reported Tango system [6, 7], where the C-terminus of a GPCR is fused with a tail sequence from AVPR2 (V2) and a transcription factor (e.g, tTa). Instead, we replaced this transcription factor with a dCas9 effector (**Fig. 1a**, left). An adaptor protein, beta2-Arrestin (ARRB2), which interacts with V2 upon GPCR activation, is fused to a <u>T</u>obacco <u>E</u>tch <u>V</u>irus protease (TEVp). To release the dCas9 effector, TEVp specifically cleaves the <u>T</u>EV <u>c</u>leavage <u>s</u>equence (TCS) placed at the N-terminus of dCas9. In the second design, we implemented a novel configuration called ChaCha, where the dCas9 effector is instead fused at the C-terminus of ARRB2 adaptor via the TCS, and the TEVp is fused at the C-terminus of GPCR via the V2 tail (**Fig. 1a**, right). In this case, ligand-activated GPCR-V2-TEVp proteolytically cleaves ARRB2-TCS-dCas9 to release the dCas9 effector, which then translocates into the nucleus to modulate genes of interest in the genome. We termed the system as ChaCha, in contrast to the Tango assay [6, 7], as both represent a functional interaction between two molecules. The Tango design generates one dCas9 effector molecule from each ligand-activated receptor. We hypothesized that the ChaCha design offers the possibility of signal amplification due to multiple effector release per ligand-receptor activation event, and thus better gene regulation performance than Tango.

**Figure 1.**
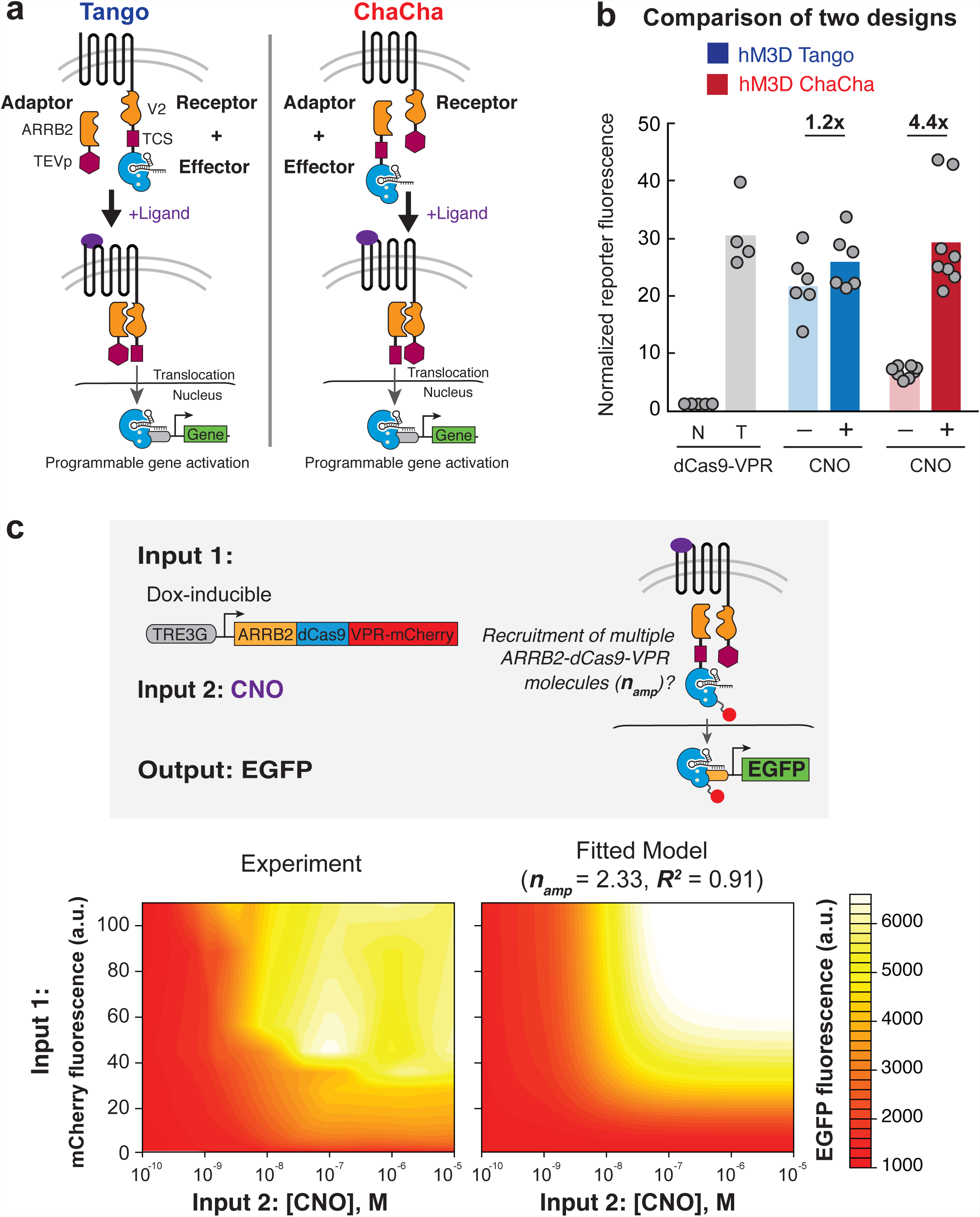
The new ChaCha design outperforms Tango. (**a**) Two design schemes of coupling CRISPR-Cas9 function to the activity of GPCRs. Left: the Tango design fused the effector protein (dCas9-VPR as shown) to the C-terminus of GPCR receptor via a V2 tail sequence. An adaptor protein, ARRB2, is fused to the TEVp protease. Right: in the ChaCha design, we fused the effector to the adaptor protein via the TCS, and fused the TEVp protease to the receptor via the V2 tail. Upon ligand binding to the receptor, both systems recruit the adaptor to the V2 tail, and the protease specifically cleaves at the TCS sequence, releasing effector proteins that translocate into the nucleus for target gene regulation. ARRB2, beta2-Arrestin; TEVp, Tobacco Etch Virus protease; TCS, TEV cleavage sequence. (**b**) Comparison of performance of unbound dCas9-VPR with Tango (blue) and ChaCha (red) using the synthetic GPCR, hM3D. N, no sgRNA; T, targeting sgRNA; +/- indicates with or without clozapine-N-oxide (CNO), the ligand for hM3D. The fold of activation displayed on top compares +/-CNO conditions. The data represent two independent experiments with technical replicates, and the bars represent the mean. *P* values from Welch’s two-sided *t*-test are provided in **Supplementary Table 1**. (**c**) Top, schematic of a stable cell line containing hM3D-CRISPR ChaCha to determine signal amplification via multiple recruitment of ARRB2-dCas9-VPR. Left, GFP activation levels after 3 days of doxycycline and CNO treatment. Right, GFP activation levels based on the rate-model fit at steady state (*R^2^* = 0.91); *n_amp_* is an index of signal amplification (see **Box 1**).

### The ChaCha design outperforms the Tango assay

Both ChaCha and Tango designs were implemented with the evolved human muscarinic 3 GPCR (hM3-DREADD or hM3D), which recognizes the bioinert small molecule, clozapine-Noxide (CNO) [12]. For gene activation, we used the *S. pyogenes* dCas9 fused to a tripartite transcriptional activator composed of VP64, p65 and Rta (dCas9-VPR) [23]. To test these constructs, we used HEK293T cells harboring a genomically integrated TRE3G promoter driving an EGFP reporter and an sgRNA targeting TRE3G. We found that the hM3D-CRISPR ChaCha design exhibited a better signal-to-noise ratio (4.4-fold activation above the ‘– CNO’ condition) compared to Tango (1.2-fold) after 1-day treatment with CNO (**Fig. 1b**). The Tango design also displayed much higher leakiness than ChaCha with significant GFP expression without CNO treatment. This may imply that fusing a large domain such as dCas9-VPR (~220 kDa, compared to 28kDa of TEVp) to a GPCR compromised receptor conformation.

### Modeling signal amplification of CRISPR ChaCha

Another explanation for the superior performance of Chacha is its potential for signal amplification. To test this hypothesis, we investigated the signal amplification feature of the ChaCha design via experiments and mathematical modeling. We formulated a set of rate equations to model the effects of ARRB2-dCas9-VPR-mCherry levels and CNO concentration on target gene (GFP) activation (**Box 1**). We generated two stable reporter HEK293T cell lines, one containing a Doxycycline (Dox)-inducibled Cas9-VPR-mCherry, and the other containing the ChaCha design with a Dox-inducible adaptor (ARRB2-dCas9-VPR-mCherry) and a constitutively expressed receptor (hM3D-V2-TEVp) (see **Methods**). Both cell lines use dCas9-VPR or released dCas9-VPR molecules to activate a genomically integrated UAS-driven GFP reporter (**Fig. 1c** and **Supplementary Fig. 1**).

As shown for the Dox-inducible dCas9-VPR cells (**Supplementary Fig. 1**), GFP expression was dependent on the expression levels of dCas9-VP R-mCherry, but not on CNO treatment. A simple DNA-binding model indicated weak power-law dependence of GFP expression to dCas9-VPR levels (power-law exponent *n* = 0.708, *R^2^* = 0.95; **Supplementary Note 1** and **Supplementary Fig. 1a**). This weak power-law dependence corroborates the observed single-turnover activity of Cas9 binding to DNA [24].

In the CRISPR ChaCha cell line, we modeled GFP expression as a hyperbolic function of both ARRB2-dCas9-VPR-mCherry and CNO ligand concentration, derived from rate equations that parsimoniously described the CRISPR ChaCha process (**Box 1**). An unbiased non-linear regression fitting of GFP expression in response to varying levels of ARRB2-dCas9-VPR and CNO (**Fig. 1c** and **Supplementary Note 2**) revealed an amplification index, ***n_amp_***, of 2.33 (*R^2^* = 0.91). The fitted amplification index for CRISPR ChaCha (***n_amp_*** > 2) implies that multiple dCas9 effector molecules are released for each ligand-activated receptor. Indeed, setting the ***n_amp_*** to 1 in the model to simulate one dCas9 effector molecule per activated receptor (e.g., Tango architecture) did not recapitulate the strong GFP response measured in our experiments (**Supplementary Fig. 1b**). Our combined experimental measurements and mathematical modeling confirmed that the Chacha design exhibits signal amplification, a feature that is critical for achieving efficient GPCR-mediated responses.

### Characterization of design parameters that impact CRISPR ChaCha performance

We characterized various design parameters that modulate the efficiency of CRISPR ChaCha, using hM3D as a model system. These parameters included (1) the linker length between ARRB2, TCS, and dCas9; (2) the proteolytic cleavage efficiency of different TCS sequences; and (3) the expression of hM3D-V2-TEVp via different promoters (**Fig. 2**). The original ChaCha design shown in **Fig. 1** (designated as variant **A**) contained a flexible linker of 3 glycine/serine repeats (GSx3) between ARRB2 and TCS. We varied the linker lengths and their locations before or after the TCS (**Fig. 2a**). Removing the linker (variant **B**) led to ~50% reduction of GFP activation upon CNO treatment, suggesting that a flexible linker facilitates hM3D-V2-TEVp access to the TCS. Adding a shorter (variant **C**, GSx1) or a longer (variant **D**, GSx5) linker before the TCS restored GFP activation, while introducing a short linker (variant **E**, GSx1) after the TCS led to leaky basal activation of GFP. The data suggested that proper placement of a flexible linker between ARRB2, TCS, and dCas9 is essential for the efficient ChaCha design.

**Figure 2.**
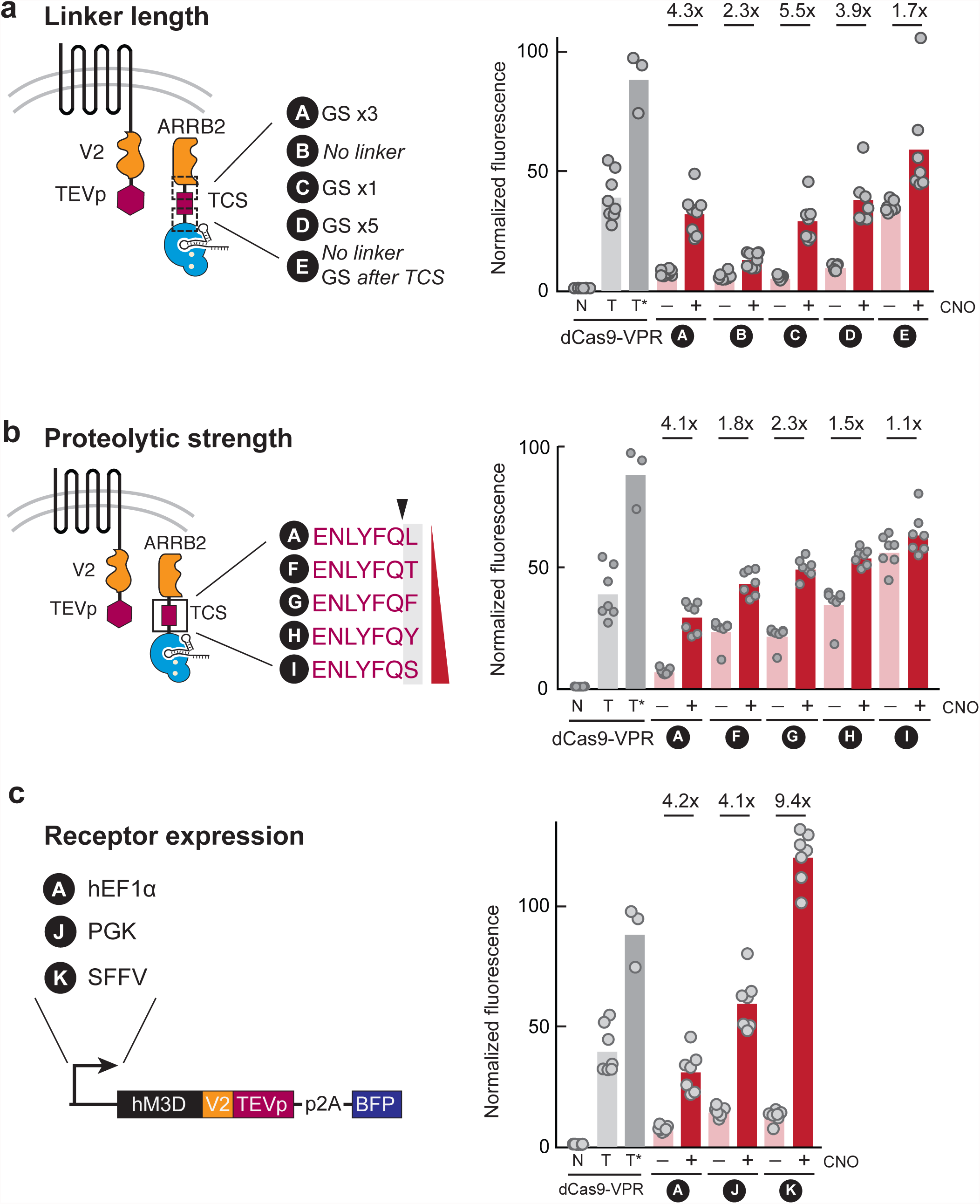
Characterizing design parameters of hM3D-CRISPR ChaCha. (**a**) Left, flexible glycine-serine (GS) linkers before or after TEV cleavage site (TCS) were varied as indicated. Right, comparison of GFP activation for variants **A** – **E**, after 1 day of +/- CNO treatment. (**b**) Left, TCS variants **F – I** with increasing proteolytic strengths. Red wedge indicates increasing proteolytic cleavage efficiency. Right, comparison of GFP activation for these variants after 1 day of +/- CNO treatment. (**c**) Comparison of GFP activation by receptor promoter variants (**A, J, K**) as indicated, after 1 day of +/- CNO treatment. N, no sgRNA; T, targeting sgRNA, 1 day posttransfection; T*, targeting sgRNA, 2 days post-transfection. +/- indicates with or without CNO. The fold of activation displayed on top compares +/-CNO conditions. The data represent three independent experiments with technical replicates, and the bars represent the mean. *P* values from Welch’s two-sided *t*-test are provided in **Supplementary Table 1**.

We then tested how proteolytic cleavage efficiency affects reporter gene activation. We modified the TCS as a spectrum of reported sequences (ENLYFQ/X, where X is any amino acid in variants **F** – **I**) [25] (**Fig. 2b**). Somewhat surprisingly, we observed that a weaker proteolytic activity achieved the highest dynamic range of the reporter gene (variant **A).** More efficient proteolytic cleavage dramatically increased the basal activation of the reporter gene in the absence of CNO, and effectively reduced the dynamic range of activation (~2-fold).

We also measured how receptor hM3D-V2-TEVp expression level affected gene activation by using different promoters. The human EF1α (hEF1α) promoter in variant **A** was replaced with PGK or SFFV promoters (variants **J** and **K)** (**Fig. 2c**). While variant **J** exhibited similar fold activation upon CNO treatment, it also exhibited ~2-fold higher basal activity without CNO. Interestingly, variant **K** exhibited the highest dynamic range of activation. Therefore, varying the stoichiometric ratio between receptor and adaptor molecules is an important consideration for optimizing the ChaCha performance in each cell type.

### Characterization of kinetics and dose response of CRISPR ChaCha

To characterize the kinetics of gene regulation in the ChaCha system, we tracked GFP activation via live-cell time-lapse microscopy (**Fig. 3a**, **Supplementary Fig. 2** and **Supplementary Movies**). We transfected the ChaCha variant **K** (**Fig. 2c**) into cells 24 h prior to CNO treatment. We saw that mCherry-tagged ARRB2-TCS-dCas9-VPR proteins were predominantly localized to the cytoplasm during 48 hours of imaging with or without CNO treatment (**Supplementary Movies 1** and **2**). GFP expression was evident as early as 12 h after addition of CNO (**Supplementary Movies 1** and **3**). In contrast, samples without CNO treatment showed little GFP activation. The observation that gene activation occurs within 12 to 24 h suggests that the ChaCha system can be used to generate fast cell responses to environmental stimuli.

**Figure 3.**
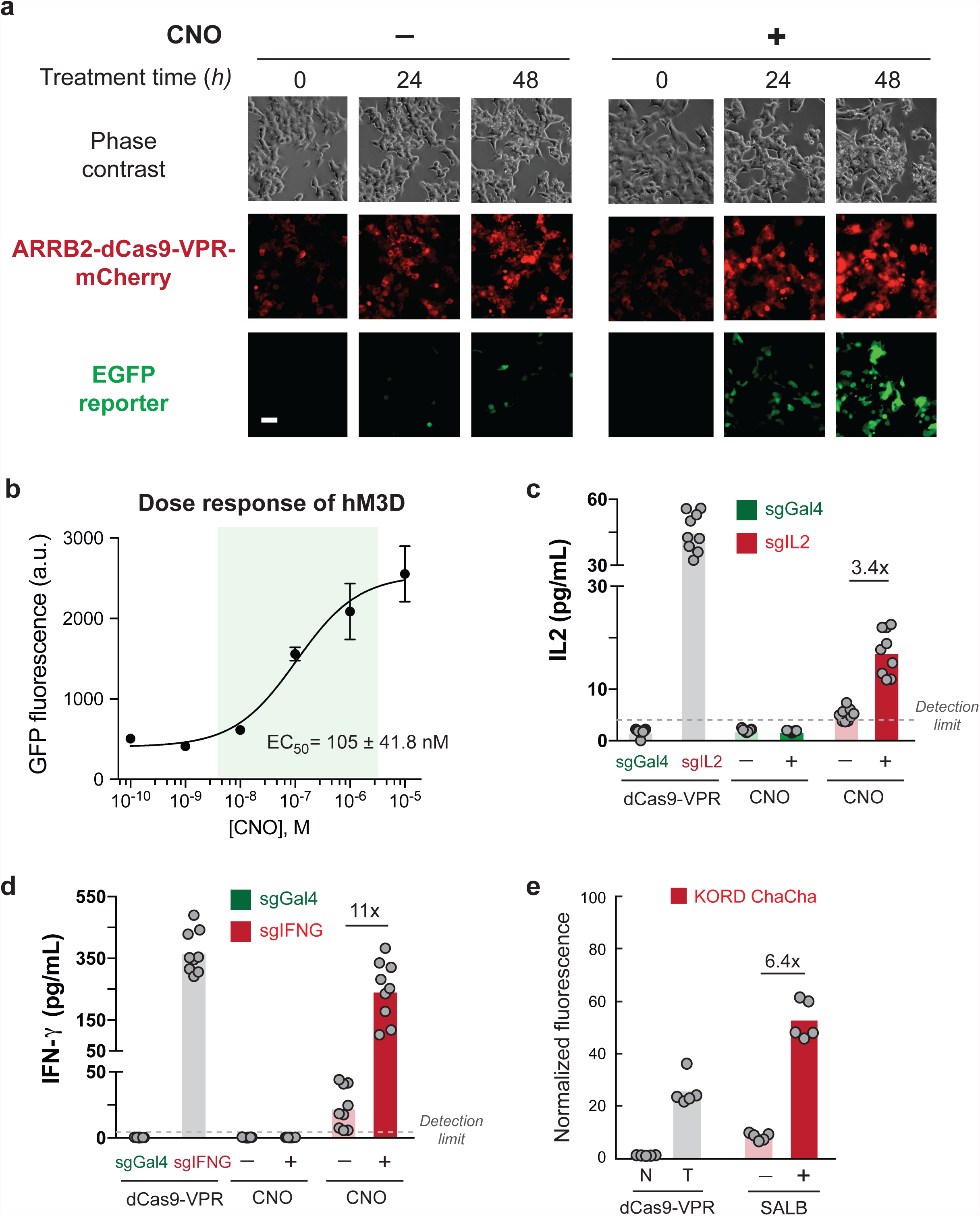
Characterization of kinetics, dose response, and input/output modularity of CRISPR ChaCha. (**a**) Time-lapse imaging of hM3D-CRISPR ChaCha in HEK293T cells over 48 hours of +/- CNO treatment. Scale bar, 50 μm. (**b**) GFP activation by hM3D-CRISPR ChaCha after 1-day treatment with increasing doses of CNO. EC50, the effective ligand concentration to achieve half-maximal GFP induction, and is shown as mean ± standard deviation (S.D.). (**c,d**) Induction of endogenous IL2 or IFN-γ expression and secretion by hM3D-CRISPR ChaCha, after 2 days of 10 μM CNO treatment. sgGal4, non-targeting sgRNA; sgIL2, *IL2*-targeting sgRNA; sgIFNG, *IFNG*-targeting sgRNA. +/- indicates with or without CNO. (**e**) Performance of KORD-CRISPR ChaCha using the GFP activation assay. N, no sgRNA; T, targeting sgRNA, 1 day post-transfection. +/- indicates with or without salvinorin B (SALB). The fold of activation displayed on top of bars compares +/- treatment conditions. The bars represent the mean. For (**b**), (**c**,**d**) and (**e**), the data represent one, three, and two independent experiments with technical replicates, respectively. *P* values from Welch’s two-sided *t*-test are provided in **Supplementary Table 1**.

We next investigated how gene activation by hM3D-CRISPR ChaCha varied with CNO ligand dose. The dose response curve for hM3D-CRISPR ChaCha displayed a relatively linear behavior, spanning over 2 orders of magnitude of ligand induction (**Fig. 3b,** and **Supplementary Fig. 3**). The fitted dose response curve based on the Hill Equation revealed the effective CNO concentration to achieve half-maximal GFP level (EC_50_) at 105 ± 4 nM (*R^2^* = 0.95, **Fig. 3b**), which is comparable to previously reported EC_50_ values using alternative methods [12].

### CRISPR ChaCha promotes efficient activation of endogenous genes

A key advantage of coupling GPCR signaling to the CRISPR-Cas9 system is Cas9’s ability to flexibly regulate the mammalian genome in a programmable manner. To demonstrate this ability in the CRISPR ChaCha system, we chose to activate two endogenous genes in HEK293T cells: interleukin 2 (*IL2*) and Interferon gamma (*IFNG*), which are cytokine genes that functionally modulate the cell-killing activity of leukocytes. IL2 and IFNG are not actively expressed in HEK293T cells (**Supplementary Fig. 4**).

**Figure 4.**
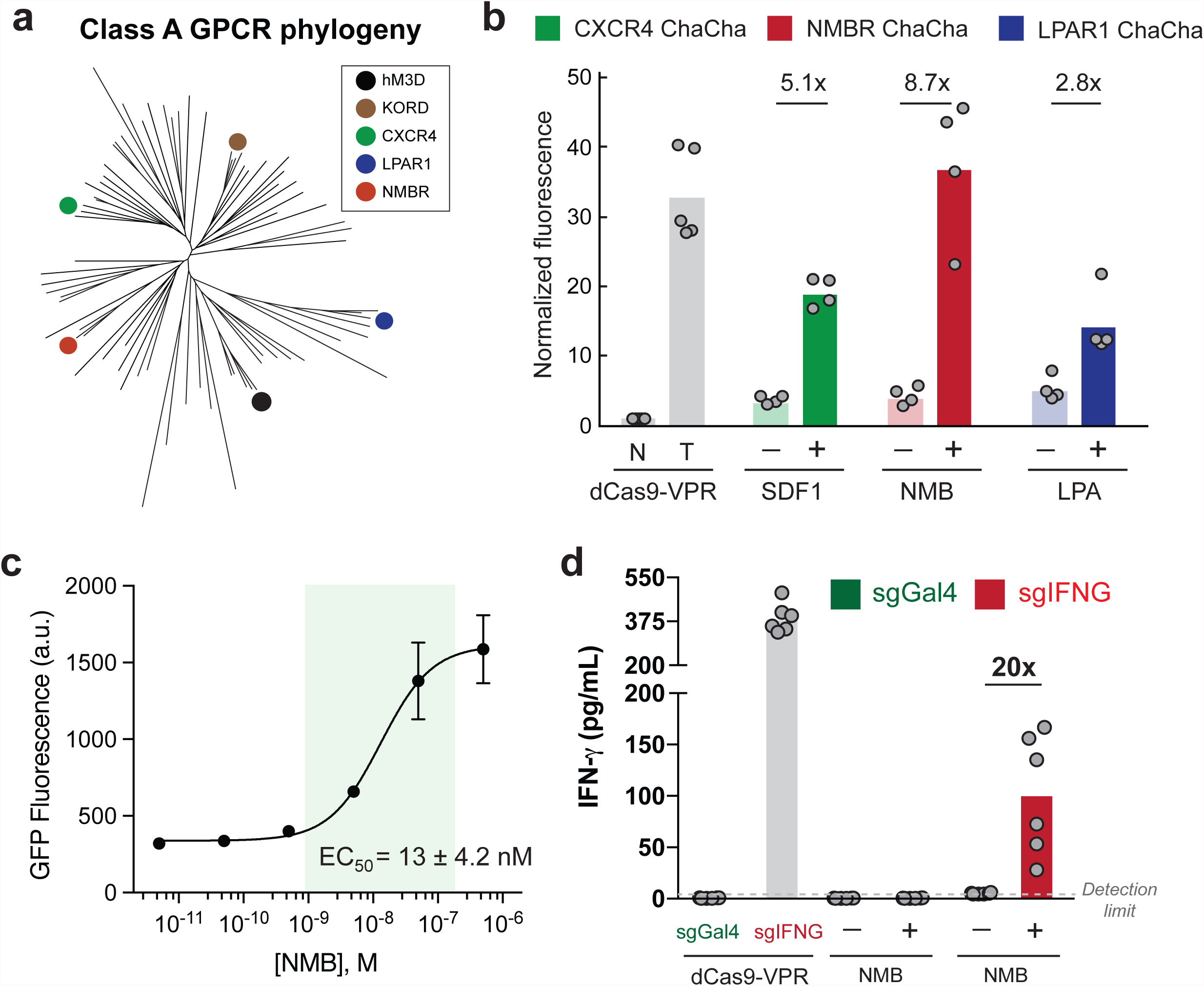
ChaCha-fying natural GPCRs. (**a**) The phylogenetic tree of Class A GPCRs. Synthetic (hM3D, KORD) and natural (CXCR4, NMBR, LPAR1) GPCRs tested here are indicated. (**b**) GFP activation of best performing ChaCha variants of CXCR4 (variant **M**), NMBR (variant **A**), LPAR1 (variant **L**) (**Supplementary Figure 5**). N, no sgRNA; T, targeting sgRNA, 1 day post-transfection; +/- indicates with or without 1-day treatment with stromal cell-derived factor 1 (SDF1, also known as CXCL12), neuromedin B (NMB), or lysophosphatidic acid (LPA) under serum-free conditions. (**c**) Representative experiment of GFP activation by NMBRCRISPR ChaCha after 1-day treatment with increasing NMB dose. EC50, the effective ligand concentration to achieve half-maximal GFP induction, and is shown as mean ± standard deviation (S.D.). (**d**) Induction of IFN-γ by NMBR-CRISPR ChaCha, after 2 days of 0.5 μM NMB treatment. sgGal4, non-targeting sgRNA; sgIFNG, *IFNG*-targeting sgRNA. +/- indicates with or without NMB. The fold of activation displayed on top of bars compares +/- treatment conditions. The bars represent the mean. For (**b,e**), and (**c**), the data represent two and one independent experiment(s) with technical replicates, respectively. *P* values from Welch’s two-sided *t*-test are provided in **Supplementary Table 1**.

We designed sgRNAs that efficiently activated each endogenous gene using the dCas9-VPR system and validated them using the appropriate Enzyme-Linked Immunosorbent Assay (ELISA) (**Supplementary Fig. 4**, see **Supplementary Methods**). We used the most effective sgRNA in combination with the hM3D-CRISPR ChaCha system for ligand-mediated endogenous gene activation. After 48-h CNO treatment, we observed efficient upregulation (3.4-fold for IL2, and 11-fold for IFN-γ) above almost undetectable levels in the absence of ligand (**Fig. 3c,d**). In contrast, the non-targeting sgRNA (sgGal4) showed no response to the CNO ligand for both genes. The data confirmed that the GPCR-coupled CRISPR system could modulate endogenous genes in human cells. The experiment above is also a proof-of-principle application of hM3D-CRISPR ChaCha as an efficient signal converter of a bioinert signal (CNO) into bioactive cytokines (IL2 and IFN-γ).

### Expanding the Chacha design to other synthetic GPCRs

We also implemented the ChaCha architecture into another evolved GPCR derived from kappa opioid receptor, KOR-DREADD or KORD, which responds to the ligand, salvinorin B (SALB) [11] (**Fig. 3e**). Like the hM3D ChaCha, the KORD ChaCha exhibited efficient GFP activation (6.4-fold) after 1-day SALB treatment, suggesting that the GPCR component of the ChaCha design is also modular.

### Expanding the ChaCha design to natural GPCRs

The G-protein coupled receptor (GPCR) family are the largest and most diverse group of membrane receptors in the human body. We further tested the modularity of the ChaCha design using three natural GPCRs: CXCR4, which senses the chemokine, stromal derived factor 1 (SDF1); NMBR, which senses the mitogen, neuromedin B (NMB); and LPAR1, which senses the fatty acid, lysophosphatidic acid (LPA). All 5 GPCRs (two synthetic and three natural GPCRs) tested here are Class A GPCRs from different branches (**Fig. 4A**). Since structural differences of each natural GPCR likely affect the performance of the ChaCha design, we further tested 3 different variants of fusing TEVp to the C-terminal tail of these natural GPCRs: direct fusion (variant **A**), partial (variant **L**) or complete replacement (variant **M**) (**Supplementary Fig. 4**).

The ChaCha design worked efficiently for all three tested natural GPCRs, with NMBR exhibiting the best activity (**Fig. 4b**). We further measured the dose-response behavior of NMBR-CRISPR ChaCha. In contrast to the hM3D receptor, the NMBR receptor displayed a narrower induction range (**Fig. 4c,** and **Supplementary Fig. 6**). The fitted dose response curve using the Hill Equation revealed an effective NMB concentration to achieve half-maximal GFP level (EC_50_) at 13 ± 4.2 nM (*R^2^* = 0.95), comparable to the reported EC_50_ values measured using alternative methods (27 nM [26], or 2 nM [7]).

We also tested whether natural GPCR-CRISPR ChaCha can efficiently activate endogenous genes. Using the NMBR-CRISPR ChaCha, we observed 20-fold upregulation of the endogenous IFN-γ expression in HEK293T cells after 2-day NMB treatment (**Fig. 4d**). Together, the CRISPR ChaCha system provides an efficient and modular platform that can employ diverse GPCRs for programmable sensing and endogenous gene regulation, which suggests that the strategy is likely generalizable to many other Class A GPCRs.

## DISCUSSION

Here we established a novel strategy to couple programmable CRISPR function with the sensing ability of diverse GPCR receptors. We demonstrated this strategy using two synthetic GPCRs and three natural GPCRs. The system is termed as ChaCha, in contrast to the Tango assay, as both depict the interaction between two molecules. In the Tango system, a transcriptional effector (e.g. tTA) is directly fused to the receptor, creating a “one-receptor, one-effector” system. In contrast, the ChaCha system fuses the transcriptional effector to the adaptor and the protease to the receptor, which turns the receptor into a multiuse enzymatic hub typified in most receptor signaling systems [1, 8].

The novel CRISPR-coupled GPCR ChaCha system presents several advantages. First, fusing a small protein TEVp to the receptor more likely preserves the conformational fidelity of the GPCR receptor. Our data showed that the hM3D ChaCha is less leaky than hM3D Tango without the CNO ligand (**Fig 1b**). Indeed, Tango-ized GPCRs suffer from constitutive activity in the absence of ligands [7]. Second, we show that our ChaCha design amplifies the input signal by possibly recruiting and releasing multiple Cas9 effectors for each receptor, while the Tango design can maximally release one effector per receptor. The rate model (**Box 1**) allowed quantification of the signal amplification behavior of the ChaCha system. Model fitting into experimental data from hM3D-CRISPR ChaCha suggested that on average 2.33 adaptor-effector molecules (ARRB2-dCas9-VPR) are recruited and released for each activated receptor. Third, the programmability of CRISPR-Cas system allows us to easily redirect the same (synthetic or natural) signal to different synthetic or natural transcriptional outputs (e.g. endogenous IL2 or IFNG). This dramatically expands previous approaches beyond the use of synthetic transcription factors (e.g., GAL4 or tTA) that can only produce synthetic outputs.

GPCRs are a large family of cell-surface receptors that respond to a variety of extracellular signals and which help regulate an incredible range of physiological functions, from sensation to growth to hormone response. Probing GPCR activity *in vivo* has been a major interest to understand their function [27]. Indeed, ~40% of drugs are developed towards targeting endogenous GPCRs [9]. The ligands detectable by GPCRs include hormones, cytokines, metabolites, small molecule compounds, and signaling peptides. However, the ligands of many orphan GPCRs are difficult to detect or undetectable via antibody-based methods [28]. Among GPCRs with known ligands, how they are dynamically regulated and activated remains a major mystery during development or neural signaling. With the fast kinetics and wider dynamic range of the ChaCha system, we envision that the GPCR research community will be able to more confidently validate novel ligands or drugs as well as profile GPCR signaling and function *in vitro* and *in vivo*.

We further envision that the GPCR-coupled CRISPR ChaCha system can be incorporated into diverse cells to create novel cell therapeutic programs. Given the diverse ligand spectrum of GPCRs and their importance in development, physiology, and disease [9], harnessing and rewiring GPCRs expands from the limited availability of antibody-based recognition domains (e.g. anti-CD19 CAR [29] or SynNotch [3]), to more diverse disease-relevant signals. For example, T cells equipped with GPCR-CRISPR ChaCha may be able to detect elevated GPCR signals in the tumor microenvironment and also execute tumoricidal functions such as upregulation of cytokines (such as IL2 and IFNG) or downregulation of checkpoint inhibitors (such as PD1). Additionally, stem cells equipped with this system can be directed to differentiate only upon ligand stimulation *in vitro*, and potentially *in vivo*. Altogether, the CRISPR ChaCha system represents a significant methodological advancement in the conversion of extracellular signals into targeted gene regulation programs.

## ACKNOWLEDGEMENTS

This work is funded by the Stanford Bioengineering Startup Funds given to L.S.Q. P.C.D.P.D. is supported by the Cancer Research Institute - Irvington Postdoctoral fellowship. N.K. is supported by the Stanford-NIST JIMB training grant, and an EDGE–STEM fellowship from the Stanford VPGE. L.S.Q. acknowledges support from the Pew Charitable Trusts and the Alfred P. Sloan Foundation.

## Box 1 I Rate Model of GPCR ChaCha

Here, we construct rate equations to model four connected processes: (*i*) conversion of inactive hM3DTEV (*R*) receptor to an activated state (*R\**) upon CNO ligand (*L*) binding, which leads to (*ii*) the cleavage of dCas9-VPR-mCherry (*C*) from ARRB2-dCas9-VPR-mCherry (*A*, referred hereafter simply as ARRB2-dCas9) that can be (*iii*) induced with doxycycline (*D*), and (*iv*) the subsequent activation of the target reporter gene, GFP (*G*), by cleaved dCas9-VPR (see Fig. 1a).

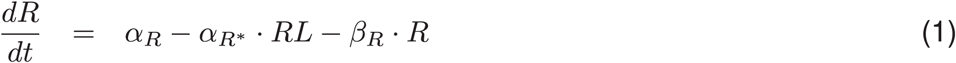

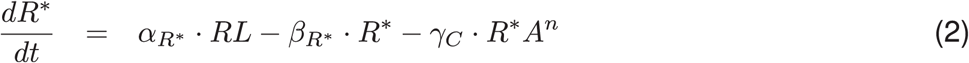

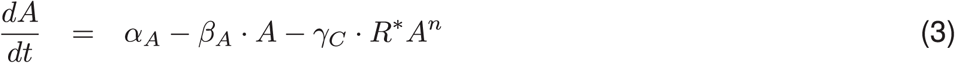

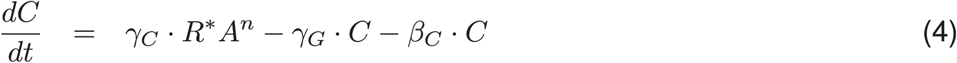

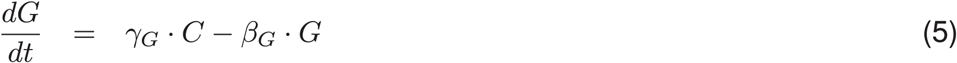

where *α_R_*, *α_R*_*, and *α_A_* are production rate constants for inactive receptor, ligand-activated receptor, and ARRB2-dCas9; *β_R_*, *β_R*_*, *β_A_*, *β_C_*, and *β_G_* are first-order degradation rate constants for inactive receptor, active receptor, ARRB2-dCas9, cleaved dCas9, and GFP, respectively; *γ_C_* and *γ_G_* are reaction rate constants for active receptor-mediated cleavage of ARRB2-dCas9 to release dCas9, and subsequent dCas9-induced production of GFP, respectively; *n* is the amplification index (i.e. *n_amp_*) representing the potential recruitment of multiple ARRB2-dCas9 molecules per one active receptor.

At steady state, all time derivatives go to zero, which yield the following steady state (*ss*) formulae for relevant molecules

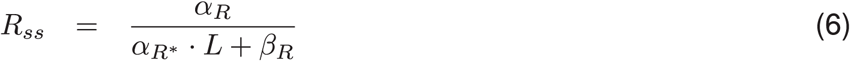

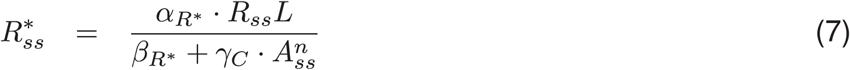

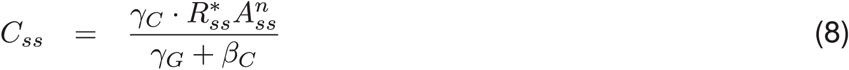

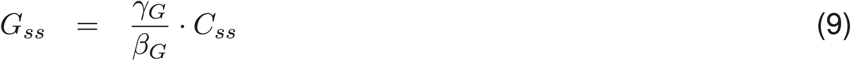

Substituting Eqs. (6), (7), (8) into Eq. (9) yields a steady-state formula for GFP as a function of CNO ligand and ARRB2-dCas9,

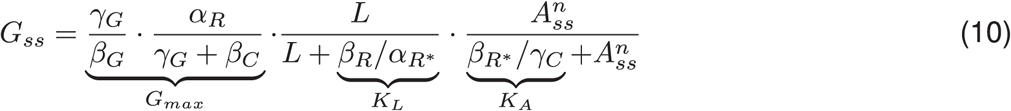

or simply,

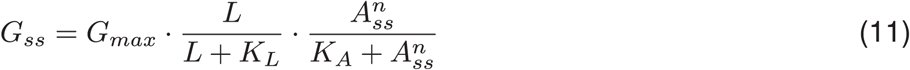

where *G_max_* represent the theoretical maximum GFP level at high saturating levels of *L* and *A*, and it is a function of the rate constants for receptor production, dCas9 degradation and GFP degradation; *K_L_* represents the set-point concentration for the CNO ligand to produce half-maximal GFP levels; *K_A_* represents the ratio of active receptor-mediated ARRB2-dCas9 cleavage and active receptor degradation rate constants.

## METHODS

The summary of statistical analyses of all experimental data is included in **Supplementary Table 1.** Standard molecular cloning techniques were performed to assemble all constructs used in this paper and they are included in **Supplementary Table 2**.

### Generation of genetic constructs

#### dCas9-effector fusions

Human codon-optimized *Streptococcus pyogenes* dCas9 was fused at the C-terminus with the tripartite VPR activator [1]. VPR is a fusion of VP64, p65 activation domains, and RTA via two GS linkers. An SV40 nuclear localization signal (NLS) was inserted C-terminal to VP64. For visualization, mCherry was fused at the C-terminus of the construct. The fusion construct was cloned into a pcDNA3 vector with a CMV promoter driving the expression of d Cas9-VPR-mCherry. For the rate model, a lentiviral pHR vector with a dox-inducible TRE3G promoter was used instead.

#### Adaptor-dCas9-effector fusions

ARRB2-TCS-dCas9-VPR was assembled by fusing ARRB2 (Human cDNA, NM_004313.3; Origene) with d Cas9-VPR-mCherry and clone into a pcDNA3 vector. The TCS sequence ENLYFQ/X was inserted in between and was flanked with GS linkers of varying lengths (**Fig. 2**). Two nuclear export signals (NES: LALKLAGLDI) flanked ARRB2 to ensure cytoplasmic localization of the chimera. For the rate model, a lentiviral pHR vector with a dox-inducible TRE3G promoter was used instead.

#### GPCR-V2-TEVp fusions

Synthetic GPCRs: hM3D (Addgene plasmid #45547) and KORD (Addgene plasmid #65417), natural GPCRs: CXCR4 (Addgene plasmid #66262), NMBR (Addgene #66445), and LPAR1 (Addgene plasmid #66418), and TEV protease (Addgene #8835) were all PCR-amplified and cloned into a pHR lentiviral vector by InFusion (Clontech) cloning. V2 sequence (derived from AVPR2) [2] was inserted in between GPCR and TEVp as primer overhangs via InFusion cloning. For visualization, p2a-BFP was fused C-terminal to TEVp. Expression of GPCR-V2-TEVp-p2a-BFP was driven by an EF1a, PGK or SFFV promoter.

All sgRNAs were cloned into a pHR lentiviral U6-driven expression vector that coexpressed puromycin-p2a-BFP or upstream of the GPCR-V2-TEVp locus for ease of transfection of the three-component GPCR CRISPR ChaCha system. Alternative sgRNA sequences were generated by PCR and inserted by InFusion cloning into the vector digested with BstXI and NotI (New England Biolabs).

### Cell culture, lentiviral production and generation of stable cell lines

HEK293T cells (Lenti-X™, Clontech) were seeded at 5×10^4^/well at day 1 in a 24-well plate format (Corning). At day 2, cells were transfected with the desired plasmids. For each well, 250 ng of each plasmid is mixed in 50 μL of Opti-MEM reduced serum media (Gibco) with 3 μL of Mirus TransIT-LT1 reagent per μg of plasmid added, and then incubated at room temperature for 15-30 minutes. Unless otherwise specified, GPCR ligands were added at the following concentrations at day 3: 20 μM CNO (Cat# C0832, Sigma), 2 μM SALB (Cat# 5611, Tocris), 0.05 μM SDF1 (Cat# 581204, BioLegend), 1 μM NMB (Cat# 1908, Tocris), and 10 μM LPA (Cat# sc-201053, Santa Cruz Biotech).

HEK293T cells were maintained in DMEM High Glucose with GlutaMAX™ media (Thermo Fisher) supplemented with 10% Tet System Approved FBS (Clontech) and 100 U/mL of penicillin and streptomycin (Gibco) at 37°C with 5% CO_2_. We did not independently authenticate these cell lines and they were not tested for mycoplasma contamination.

HEK293T cells were also used for lentiviral packaging. At day 1, cells were seeded at 2.0 – 3.0×10^5^ cells/mL in a 6-well plate format (Corning). At day 2, cells were 50-70% confluent at the time of transfection. For each well, 1.51 μg of pHR vector containing the construct of interest, 1.32 μg of dR8.91 and 165 ng of pMD2.G (Addgene) were mixed in 250 μL of Opti-MEM reduced serum media (Gibco) with 7.5 μL of Mirus TransIT-LT1 reagent and incubated at room temperature for 15-30 minutes. The transfection complex solution was distributed evenly to HEK293T cultures dropwise. Media was replaced at day 3 with fresh media. At day 4, lentiviruses are harvested from the supernatant with a sterile syringe and filtered through a 0.45-μm polyvinylidene fluoride filter (Millipore) for immediate transduction of target cell cultures.

Filtered lentiviral supernatants were mixed 1:1 with appropriate fresh media to replace media of target cells for transduction. Adherent cell cultures were transduced at 50% confluence. Approximately 10 days after transduction, the HEK293T pTRE3G-EGFP line and the pUAS-EGFP::pEF1a-rtTA-p2a-puro reporter line (pre-selected for 2 days with 1 μg/μL puromycin) were transiently transfected with dCas9-VPR and a targeting sgRNA (sgTET and sgUAS, respectively) for 1 day prior to sorting via EGFP FACS in Carmen (BD InFlux) and Aida (BD Aria II) sorters, respectively. For the rate model, the HEK293T pUAS-GFP line was transduced with pEF1a-hM3D-V2-TEVp-p2a-BFP and pTRE3G-(ARRB2-TCS)-dCas9-VPR-mCherry and sorted approximately 7 days after transduction for both BFP and 1-day doxycycline induction of mCherry expression.

### Flow cytometry analysis

Cell were dissociated using 0.05% Trypsin-EDTA (Life Technologies) and analyzed for reporter fluorescence in the Stanford Shared FACS facility with a Scanford FACScan analyzer (Becton Dickinson). For the rate model, cells were filtered into a 96-well plate for high-throughput analysis using the CytoFLEX S flow cytometer (Beckman Coulter).

### Time-Lapse Microscopy

At day 0, HEK293T TRE3G-EGFP reporter cells were plated at 1x10^5^ cells per 24-well well (μ-Plate 24 well; ibidi). At day 1, 250 ng of each plasmid was transfected (see **Fig. 3a** and **Supplementary Fig. 2**). At day 2, 20 μM of CNO was added to appropriate wells and immediately imaged (Model DMi8, Lumencor SOLA SMII 405, Leica DFC9000; Leica Microsystems) at 37 °C with 5% CO_2_. Leica Application Software was used to set up time-lapse imaging. Images from phase contrast, mCherry and GFP channels were taken every 0.5 h for 48 hours with a 20x/0.40 objective using Leica Adaptive Focus control.

### Endogenous Cytokine Activation and Secretion Assays

A day before transfection, HEK293T cells were seeded in 24-well plates at a density of 5x10^4^ cells per well. On day 1, cells were transfected with 250 ng of each plasmid (i.e. the CRISPR ChaCha components: GPCR-V2-TEVp of interest, the ARRB2-TCS-dCas9-VPR, and an sgRNA). On day 2, controls were transfected, consisting of the GPCR of interest, dCas9-VPR, and an sgRNA. Media on the ChaCha containing cells was then changed to those with or without ligand treatment (10 μM for CNO; 0.5 μM for NMB).

Supernatants from cell cultures were harvested on day 4, and stored at -80C. Secreted proteins were quantified using the ELISA MAX Deluxe kits for human IL-2 and IFN-γ (BioLegend). Absorbance at 450 nm and 570 nm was measured for samples in technical triplicates with a Synergy H1 plate reader (BioTek). Samples were standardized by subtracting measurements at 570 nm from those at 450 nm. Protein concentrations were then determined by standard curves fitted to a power law using Excel (Microsoft).

### Class A GPCR Phylogenetic Tree Construction

The phylogenetic tree in **Fig. 4a** was constructed using GPCRdb [3, 4]. Human GPCRs from the Swiss-Prot database were used as reference, without any selection for G protein preference. For GPCRs used in this study their entire family was selected, otherwise one GPCR from each family was used to construct the tree. Full-length sequences of receptors were considered for tree construction. No bootstrapping was performed, and distance calculation utilized the neighbor-joining method, utilizing the regular branch lengths option. The tree was then rendered by taking the generated newick file and viewing it in T-REX [5].

### Data Presentation and Analyses

Data are displayed as individual points with sample size indicated in figure legends. No sample size estimates were performed, and the sample sizes used in this study are consistent with those used by similar genome editing and gene regulation studies. Experiments were performed independently at least two times. Values reported are relative to indicated control conditions. No randomization or blinding was performed.

Statistical analysis was performed using SPSS Statistics 21 (version 22, IBM Corporation). Equal variance between populations was not assumed. To account for unequal variance among conditions, Welch’s two-sided *t*-test was performed when comparing two conditions, and Welch’s ANOVA was performed followed by Games–Howell *post hoc* tests when comparing more than two conditions with each other. All statistical data analyses are compiled in **Supplementary Table 1**.

## Supplementary Figure Legends

**Supplementary Figure 1.**
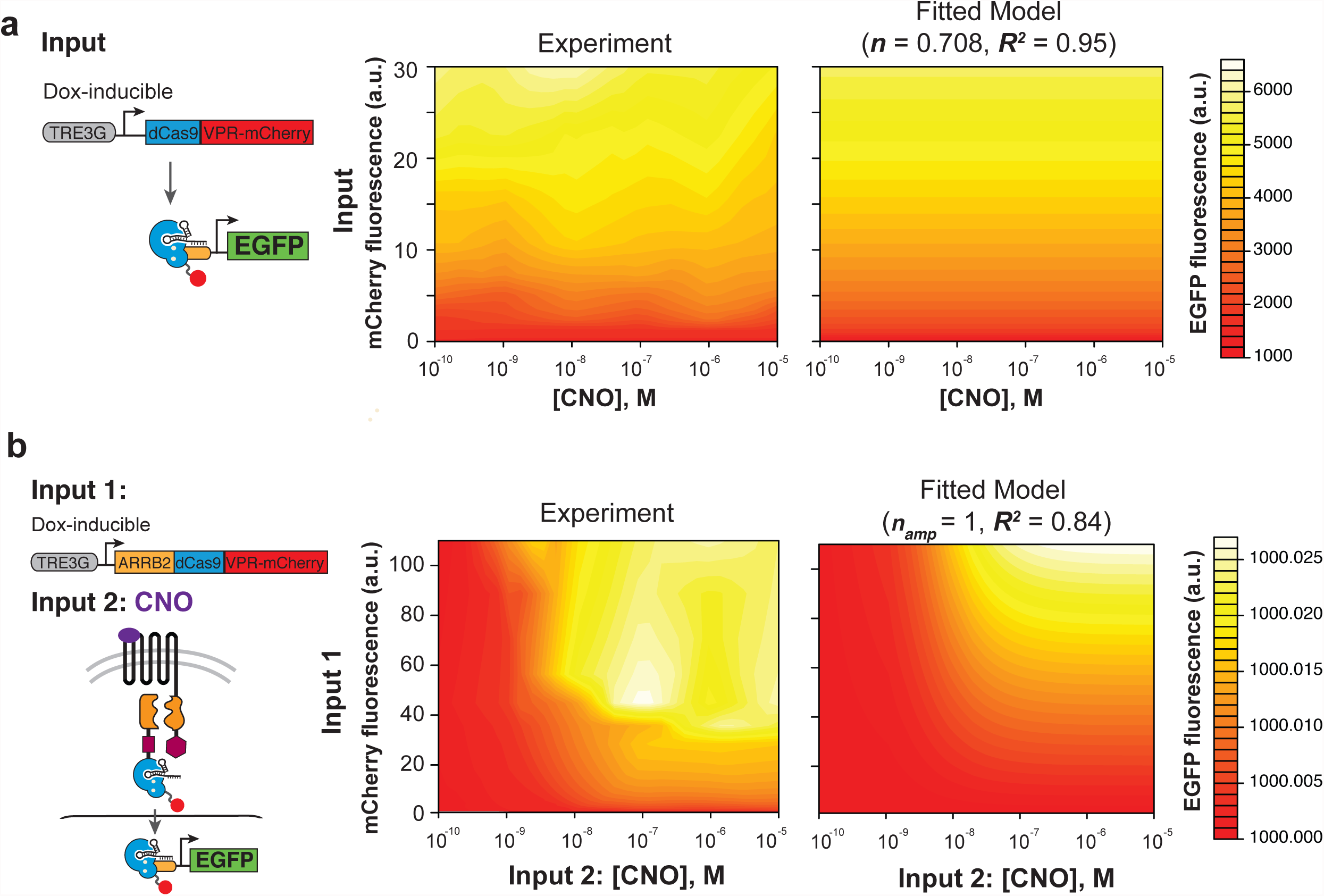
Measurement and modeling EGFP reporter activation via doxycycline-inducible dCas9-VP R-mCherry or doxycycline-inducible hM3D-CRISPR ChaCha. (**a**) Left, schematic of the HEK293T stable line containing pTR E3 G-dCas9-VPR-mCherry, pUAS-EGFP::pEF1 a-rtTA-p2a-puro, and pU6-UAS-targeting sgRNA (sgUAS8). Doxycycline (Dox) induces expression of dCas9-VPR-mCherry, which with sgUAS8 binds and activates the *EGFP* reporter. Plots demonstrate GFP fluorescence (a.u) as a function of mCherry fluorescence (a.u, y-axis; a surrogate for Dox), and CNO concentration (M, x-axis). Both the experimental (middle) and fitted-model plots (right) demonstrate a Dox-dependent, CNO-independent graded EGFP response. Model fitting using non-linear regression indicates ***n_amp_*** < 1 (right). (**b**) Related to **Fig. 1c**. Left, schematic of HEK293T stable line with pTRE3G-ARRB2-dCas9-VPR-mCherry, pUAS-EGFP::pEF1a-rtTA-p2a-puro, and pU6-sgUAS8::pEF1a-hM3DTEVp. Dox induces expression of ARRB2-d Cas9-VP R-m Cherry, while hM3D-TEVp is constitutively expressed, but only activated by CNO. Both CNO and Dox are needed for reporter activation. Right, setting ***n_amp_*** to one in order to mimic the Tango design significantly reduces the variable range in which maximum gene activation is achievable. Experimental result for ChaCha is shown here for comparison (middle). All experimental results for model fitting were collected after 3 days of CNO and Dox treatment, at which point steady-state expression of EGFP is assumed.

**Supplementary Figure 2.**
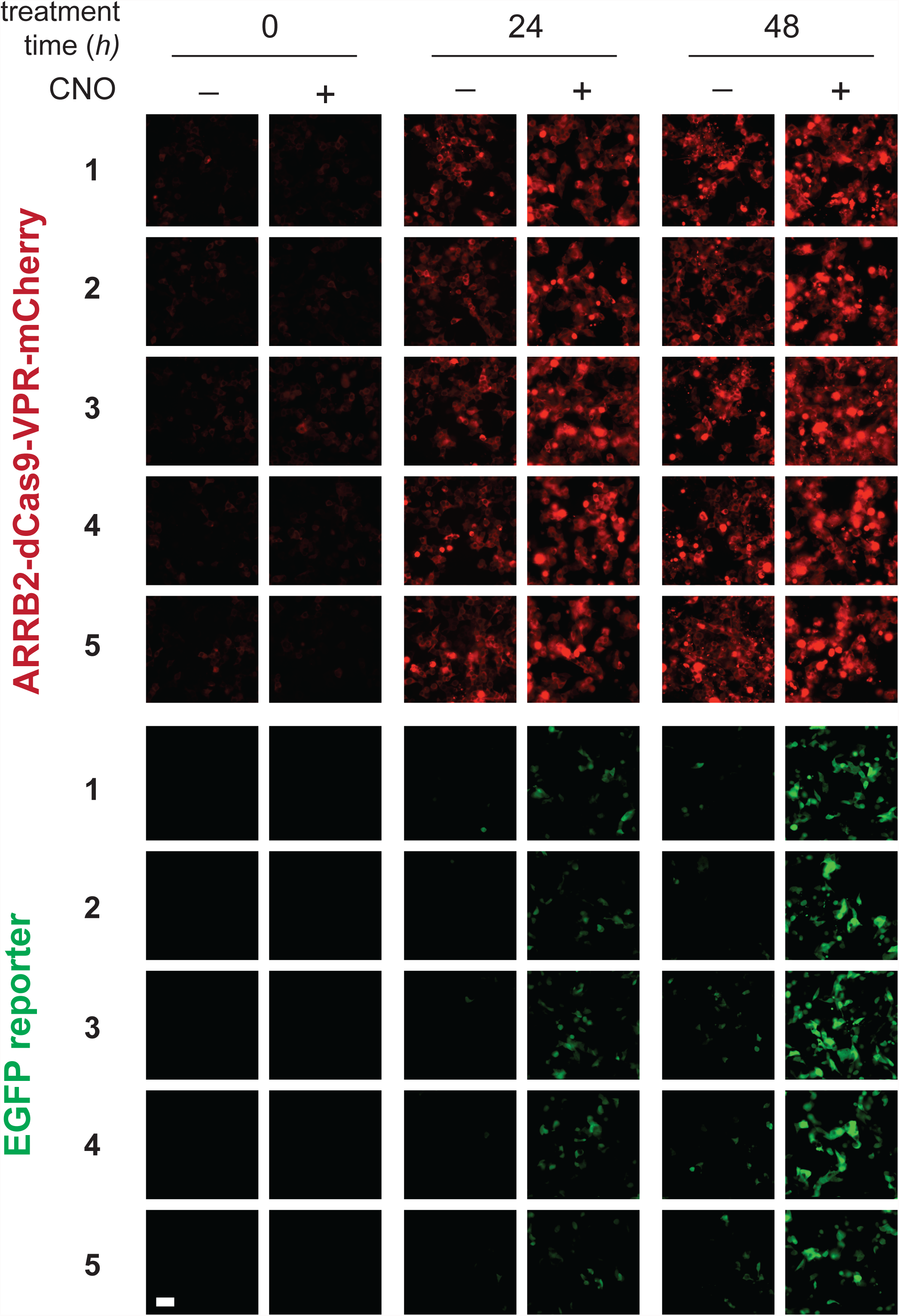
Additional fields of view from time-lapse microscopy of hM3D-CRISPR ChaCha activation in live cells. Related to **Fig. 3a** and **Supplementary Movies**.

**Supplementary Figure 3.**
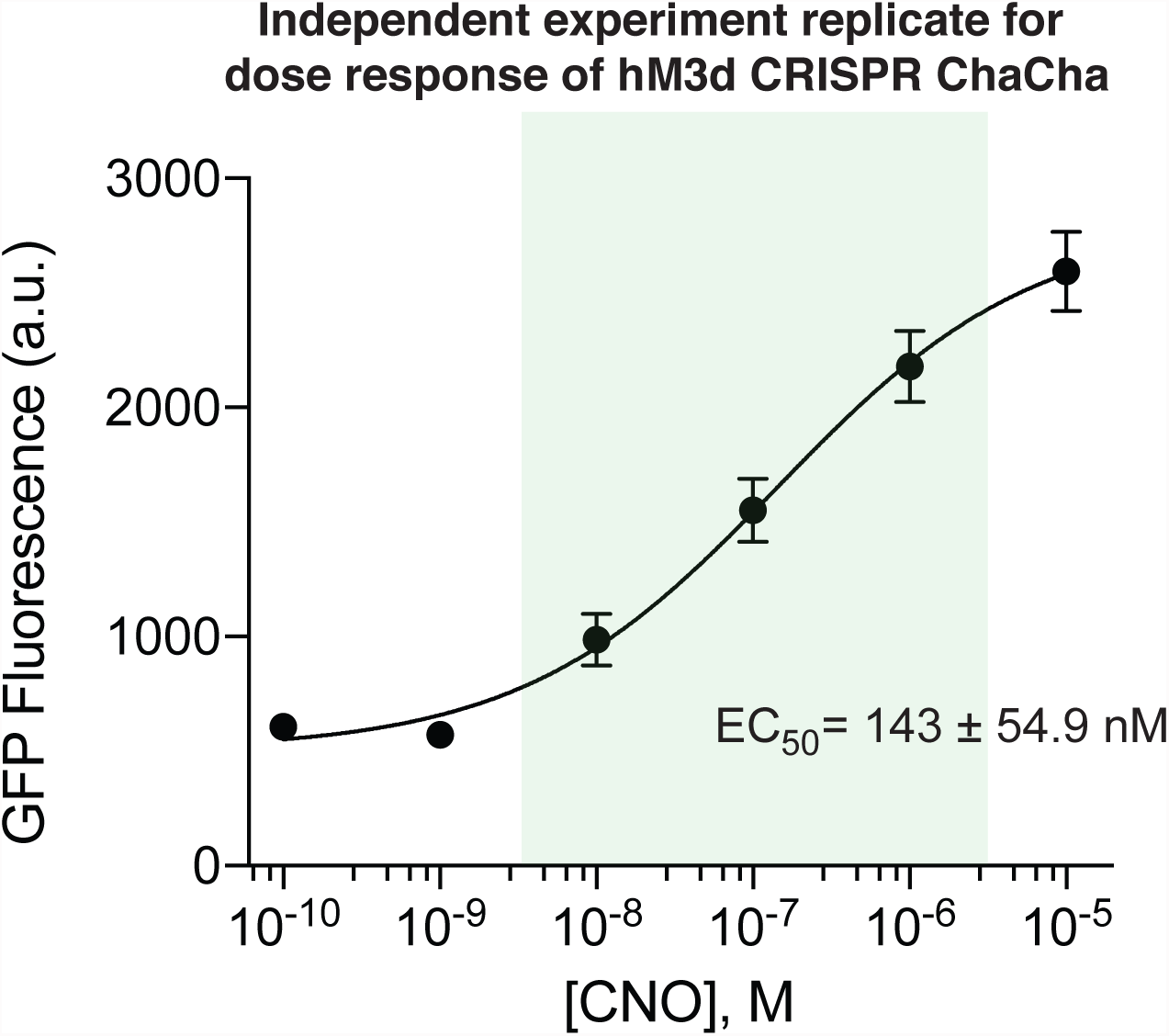
Independent Replicate of CNO Dose Response Curve for hM3D-CRISPR ChaCha. Related to **Fig. 3b**.

**Supplementary Figure 4.**
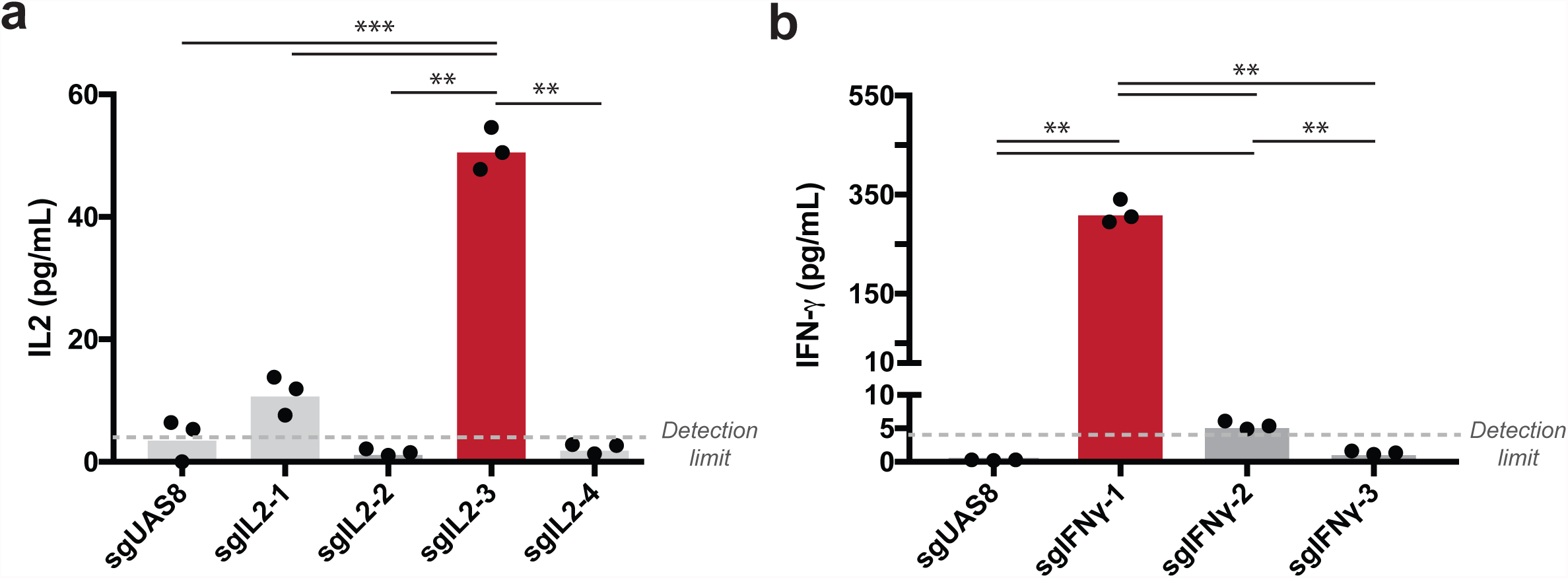
Screen of sgRNAs for IL2 and IFNG cytokine activation and secretion. HEK293T cells were transfected with dCas9-VPR and individual sgRNAs to be tested. Cytokines were measured by ELISA. Best-performing sgRNAs from each were used for experiments in the main text (n = 3 technical replicates from different wells of a 24-well plate, from one experiment). (*P < 0.05, **P < 0.01, ***P < 0.001; Welch’s ANOVA with Games-Howell post-hoc test).

**Supplementary Figure 5.**
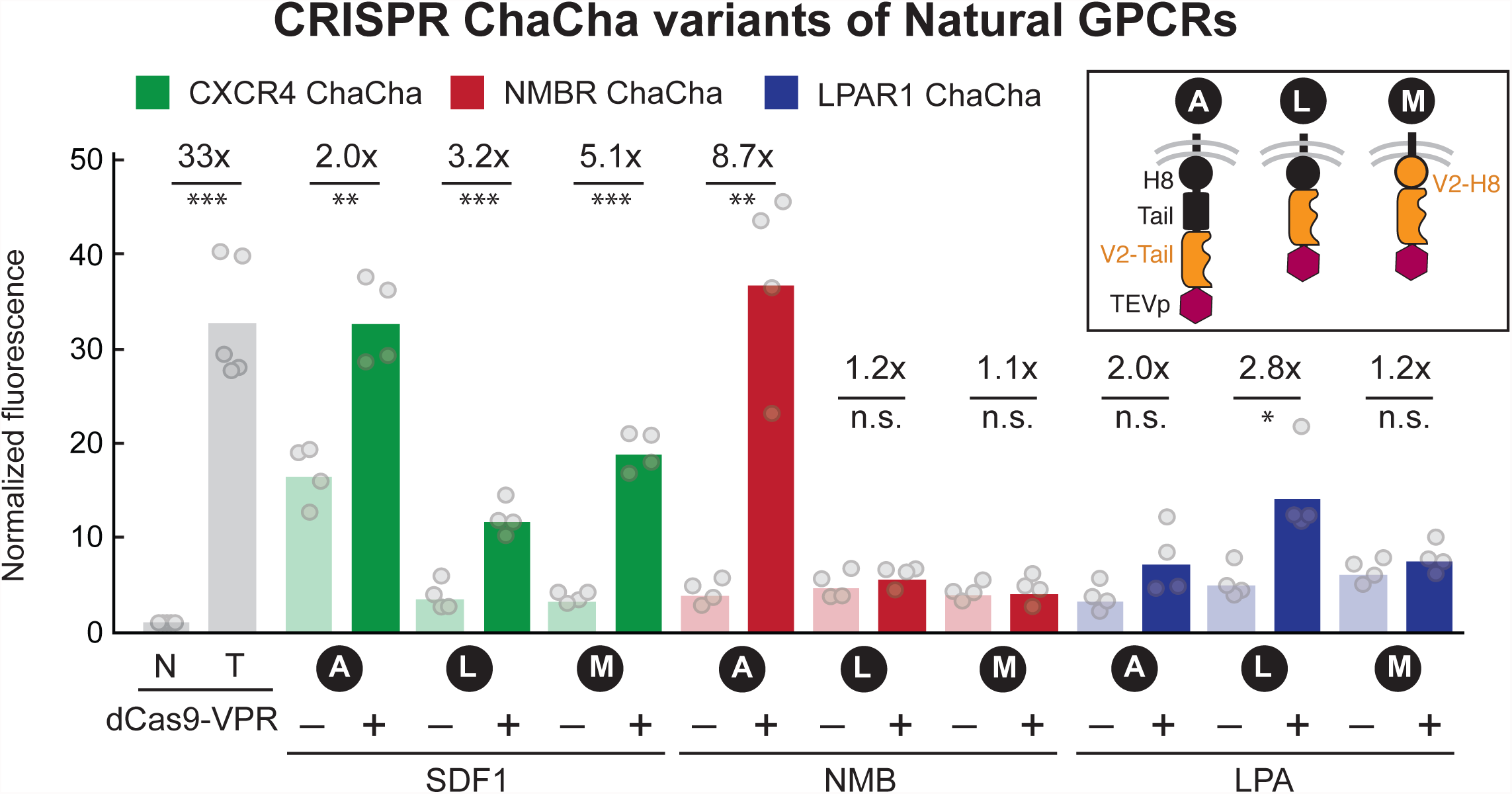
CRISPR ChaCha C-terminal tail variants of Natural GPCRs. HEK293T cells were transfected with C-terminal tail variants of the ChaCha system. Variant **A** is the principal ChaCha architecture; variant **L** is a C-terminal tail replacement with that of V2, and variant **M** is a C-terminal tail and H8 domain replacement to V2 (inset). System behavior was assayed based on GFP reporter activation with the presence (+) or absence (-) of cognate ligands for 1 day under serum-free conditions, as measured by flow cytometry. Dots represent biological measurements, and bars represent means from 2 independent experiments. The fold of activation displayed on top of bars compares +/- treatment conditions. N, no sgRNA; T, targeting sgRNA, 1 day post-transfection. (*P < 0.05, **P < 0.01, ***P < 0.001, n.s. = not significant; Welch’s two-sided t-test).

**Supplementary Figure 6.**
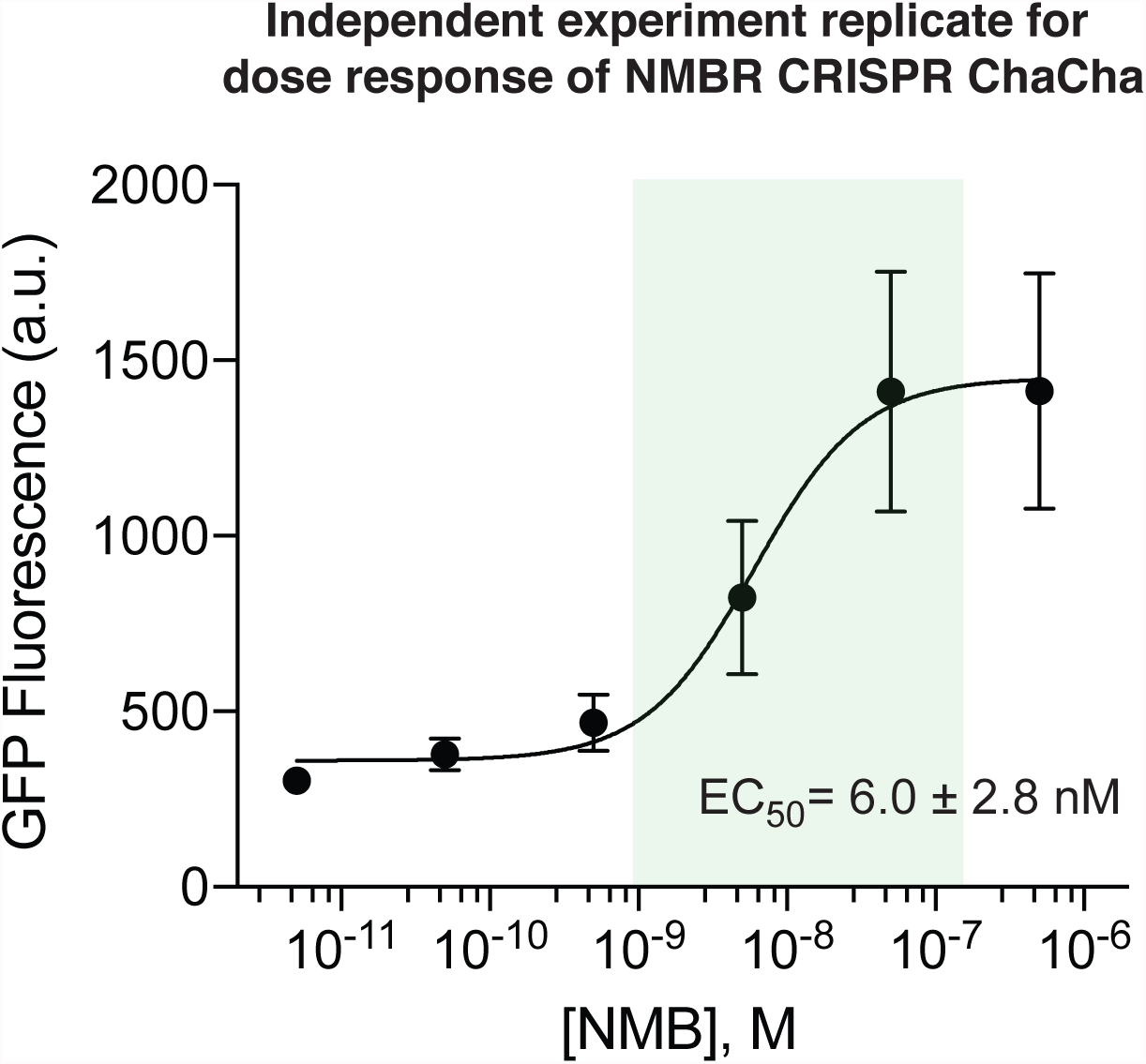
Independent Replicate of NMB Dose Response Curve for NMBR-CRISPR ChaCha. Related to **Fig. 4b**.

**Supplementary Table 1.** Statistical Analyses of Experimental Data. See figure legends and **Methods** for descriptions of statistical tests used. Fold changes are mean values that are reflected in the figures. Statistically significant differences (P < 0.05) are **bolded.**

| Figure | Condition 1 | Condition 2 | Fold | P-value |
| --- | --- | --- | --- | --- |
| **1b** | dCas9-VPR (N) | dCas9-VPR (T) | 31 | **0.002** |
|  | hM3D Tango (- CNO) | hM3D Tango (+ CNO) | 1.2 | 0.216 |
|  | hM3D ChaCha (- CNO) | hM3D ChaCha (+ CNO) | 4.4 | **<0.001** |
| **2a** | dCas9-VPR (N) | dCas9-VPR (T) | 39 | **<0.001** |
|  | dCas9-VPR (N) | dCas9-VPR (T*) | 89 | **0.007** |
|  | Variant **A** (-CNO) | Variant **A** (+CNO) | 4.3 | **<0.001** |
|  | Variant **B** (-CNO) | Variant **B** (+CNO) | 2.3 | **<0.001** |
|  | Variant **C** (-CNO) | Variant **C** (+CNO) | 5.5 | **<0.001** |
|  | Variant **D** (-CNO) | Variant **D** (+CNO) | 3.9 | **<0.001** |
|  | Variant **E** (-CNO) | Variant **E** (+CNO) | 1.7 | **0.028** |
| **2b** | dCas9-VPR (N) | dCas9-VPR (T) | 40 | **<0.001** |
|  | dCas9-VPR (N) | dCas9-VPR (T*) | 89 | **0.007** |
|  | Variant **A** (-CNO) | Variant **A** (+CNO) | 4.1 | **<0.001** |
|  | Variant **F** (-CNO) | Variant **F** (+CNO) | 1.8 | **<0.001** |
|  | Variant **G** (-CNO) | Variant **G** (+CNO) | 2.3 | **<0.001** |
|  | Variant **H** (-CNO) | Variant **H** (+CNO) | 1.5 | **<0.001** |
|  | Variant **I** (-CNO) | Variant **I** (+CNO) | 1.1 | 0.112 |
| **2c** | dCas9-VPR (N) | dCas9-VPR (T) | 40 | **<0.001** |
|  | dCas9-VPR (N) | dCas9-VPR (T*) | 89 | **0.007** |
|  | Variant **A** (-CNO) | Variant **A** (+CNO) | 4.2 | **<0.001** |
|  | Variant **J** (-CNO) | Variant **J** (+CNO) | 4.1 | **<0.001** |
|  | Variant **K** (-CNO) | Variant **K** (+CNO) | 9.4 | **<0.001** |
| **3c** | dCas9-VPR (N) | dCas9-VPR (T) | 26 | **<0.001** |
|  | dCas9-VPR (N) | dCas9-VPR (T*) | 87 | **<0.001** |
|  | KORD ChaCha (-SALB) | KORD ChaCha (+SALB) | 6.4 | **<0.001** |
| **3d** | dCas9-VPR (sgGal4) | dCas9-VPR (sgIL2) | >11* | **<0.001** |
|  | hM3D ChaCha (sgGal4, - CNO) | hM3D ChaCha (sgGal4, + CNO) | b.d.l.* | - |
|  | hM3D ChaCha (sgIL2, - CNO) | hM3D ChaCha (sgIL2, + CNO) | 3.4 | **<0.001** |
| **3e** | dCas9-VPR (sgGal4) | dCas9-VPR (sgIFNG) | >92* | **<0.001** |
|  | hM3D ChaCha (sgGal4, - CNO) | hM3D ChaCha (sgGal4, + CNO) | b.d.l.* | - |
|  | hM3D ChaCha (sgIFNG, - CNO) | hM3D ChaCha (sgIFNG, + CNO) | 11 | **<0.001** |
| **4b** | dCas9-VPR (N) | dCas9-VPR (T) | 33 | **<0.001** |
|  | CXCR4 ChaCha (- SDF1) | CXCR4 ChaCha (+ SDF1) | 5.1 | **<0.001** |
|  | NMBR ChaCha (- NMB) | hM3D ChaCha (+ NMB) | 8.7 | **0.007** |
|  | LPAR1 ChaCha (- LPA) | hM3D ChaCha (+ LPA) | 2.8 | **0.024** |
| **4d** | dCas9-VPR (sgGal4) | dCas9-VPR (sgIFNG) | >97 | **<0.001** |
|  | NMBR ChaCha (sgGal4, - NMB) | NMBR ChaCha (sgGal4, + NMB) | n.d.* | - |
|  | NMBR ChaCha (sgIFNG, - NMB) | NMBR ChaCha (sgIFNG, + NMB) | 20 | **0.009** |

\*For ELISA values below the detection limit of 4 pg/mL, fold changes are estimates compared to that limit; b.d.l., below detectable limit.

**Supplementary Table 2.**
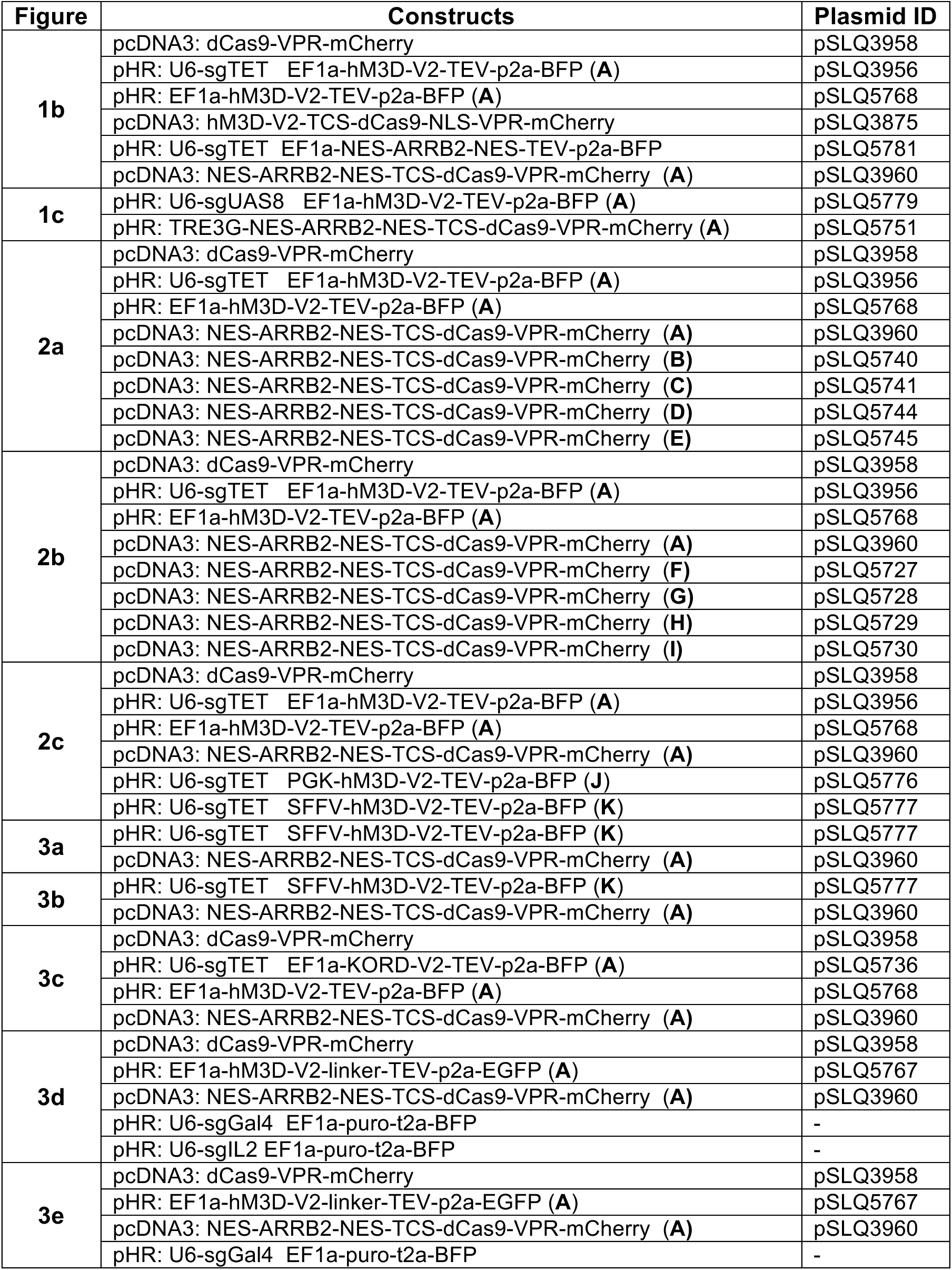
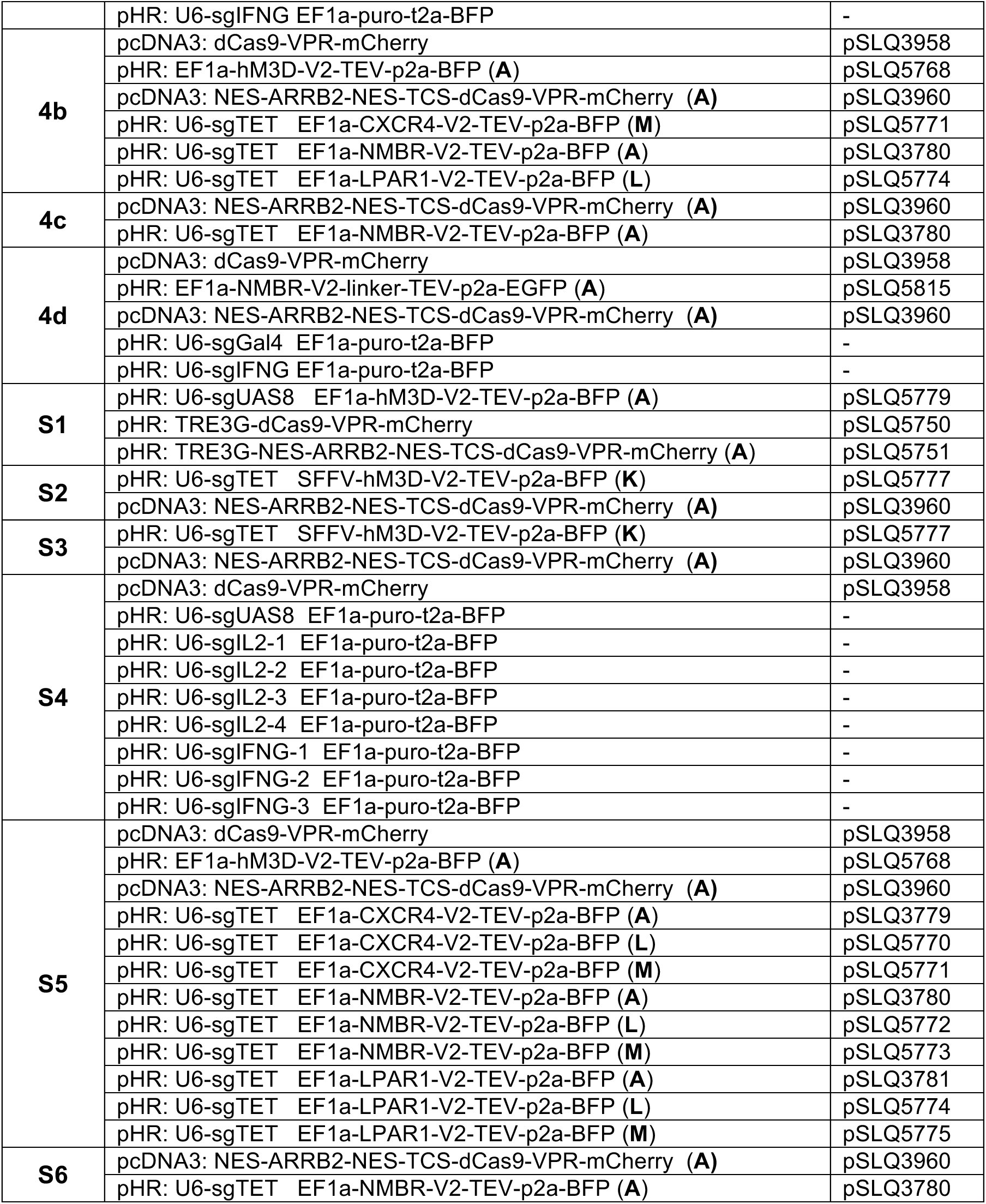
Plasmid constructs used in this study listed by figure. ChaCha variants are indicated in parentheses. Constructs used to generate the two reporter cell lines are described in the **Methods.**

## Supplementary Note 1

### Modeling GFP Activation by Dox-inducible dCas9-VPR

We construct rate equations to model the induction of dCas9-VPR-mCherry (referred hereafter simply as dCas9) by doxycycline (dox, *D*) and the dCas9-induced activation of the target reporter gene, GFP.

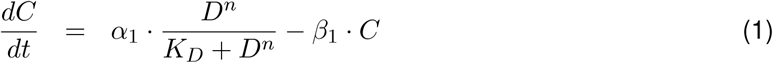

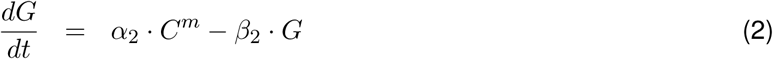

where *α*_1_ and *α*_2_ are first-order rate constants for dox-induced dCas9 (*C*) production and subsequent dCas9-induced production of GFP (*G*), respectively; the Hill coefficient *n* and *K_D_* are the cooperativity and affinity constants of dox induction, respectively; the exponent *m* is a lumped parameter that captures the following processes in series: dCas9 binding to the gene target (*GFP*), transcription, and translation of GFP; *β*_1_ and *β*_2_ are first-order degradation rate constants for dCas9 and GFP, respectively.

At steady state,

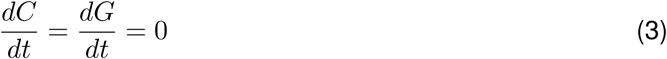

which yields steady state (*ss*) formulae for *C* and *G*

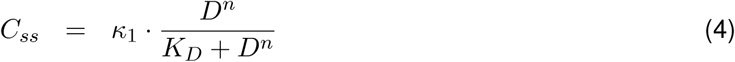

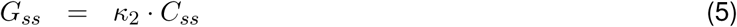

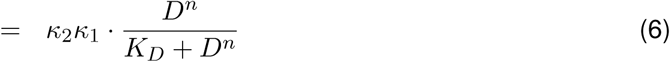

where 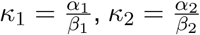, and *G_max_* = *K*_1_*K*_2_; *G_max_* represents the theoretical maximum GFP level.

## Supplementary Note 2

### Nonlinear Regression Fitting of Rate Model to Experiment

We used the open source software, RStudio (version 1.0.136), to generate plots in **Fig. 1c** and **Supplementary Fig. 1** and to determine the goodness of fit of the rate model equations to experimental data.

The purpose of the following R code is to load, interpolate and plot experimental data on GFP activation by hM3D-CRISPR ChaCha (**Fig. 1c, left** and **Supplementary Data 1 and 2**). Nonlinear regression fitting is also performed in this code to fit Equation (11) in **Box 1** to experimental data and determine parameter values and the goodness of fit.

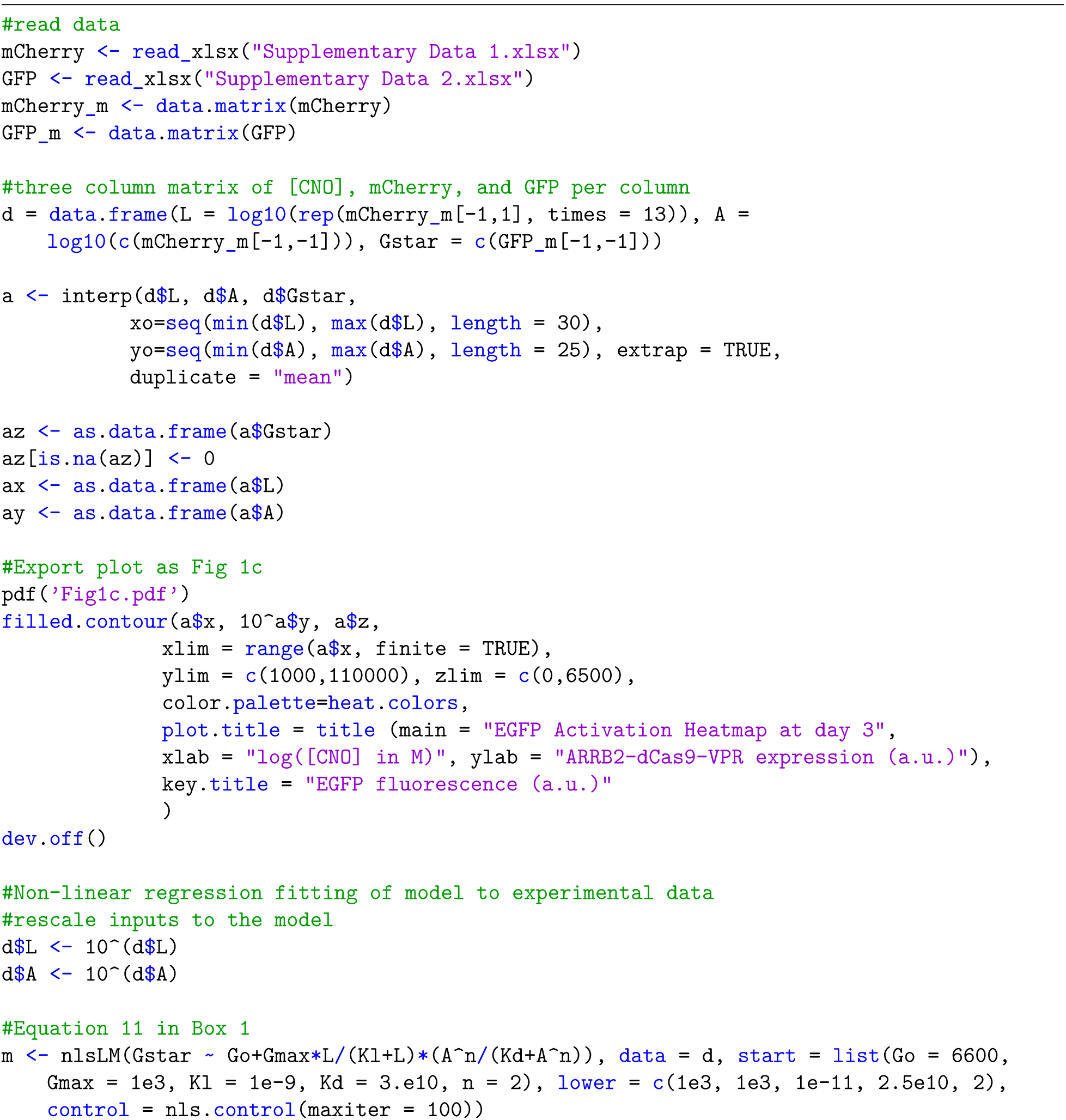
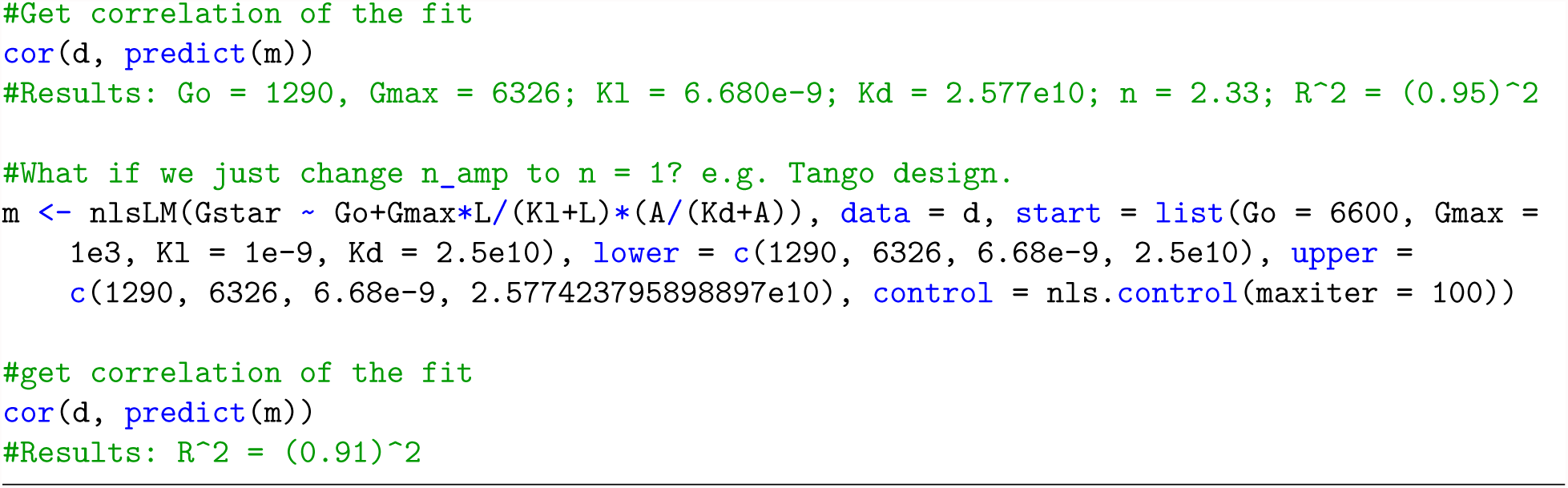

The purpose of the following R code is to plot Equation (11) in **Box 1** with the fitted parameter values (**Fig. 1c, right** and **Fig. S1b, right**).

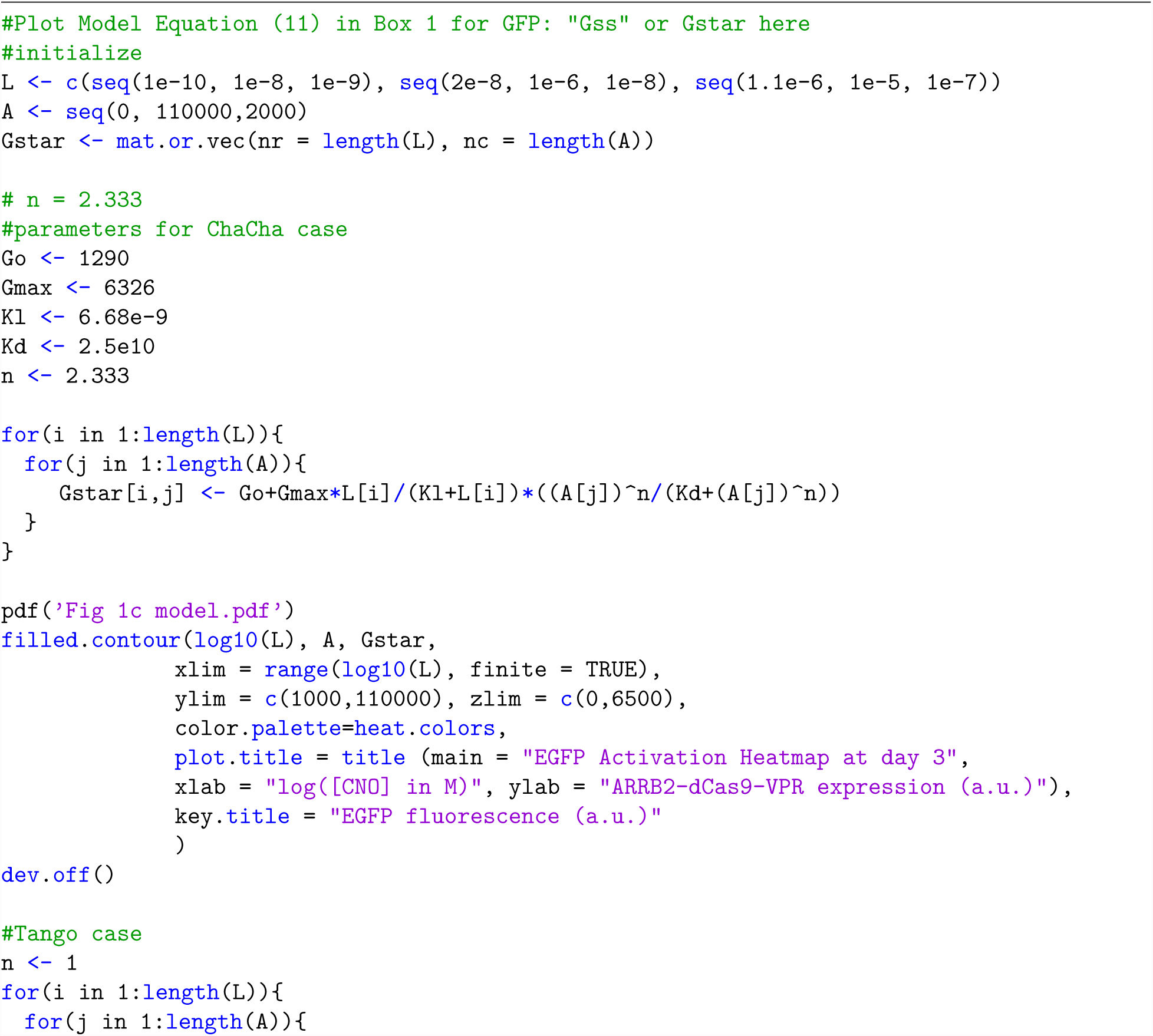
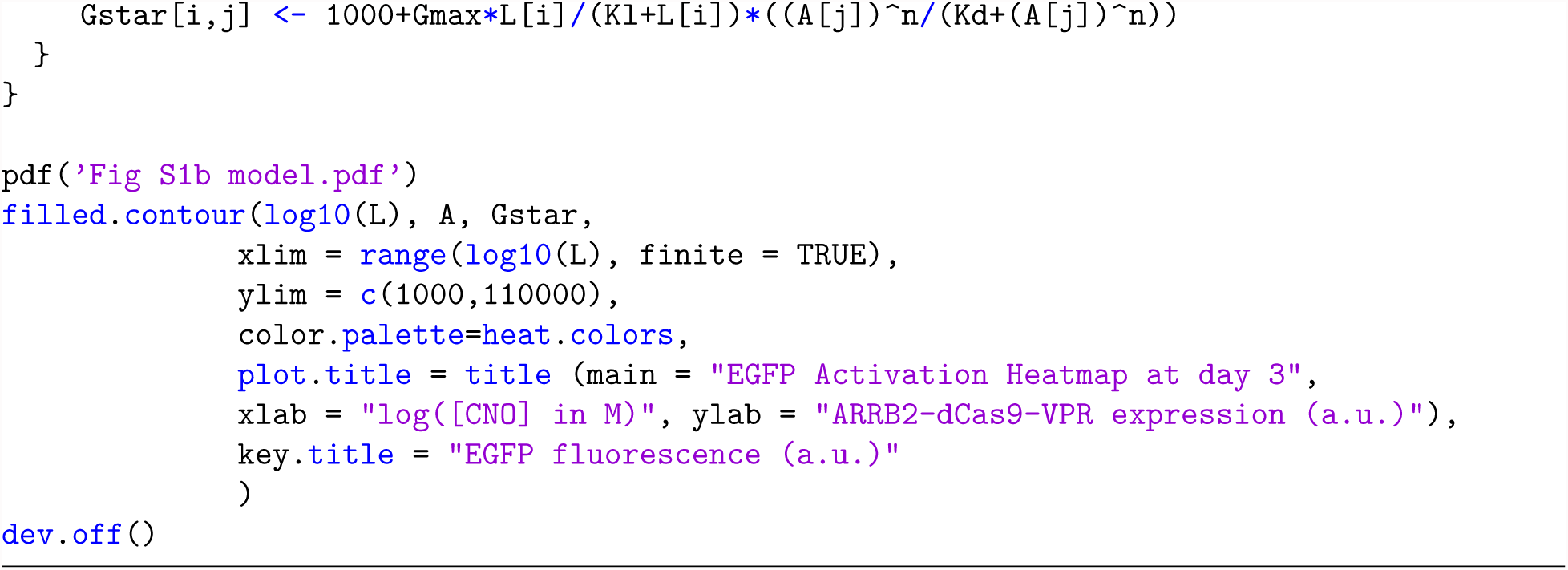

The purpose of the following R code is to load, interpolate and plot experimental data on GFP activation by dox-inducible dCas9-VPR (**Fig. S1a, left**, and **Supplementary Data 3 and 4**). Nonlinear regression fitting is also performed in this code to fit Equation (6) in **Supplementary Note 2** to the data and determine parameter values and the goodness of fit.

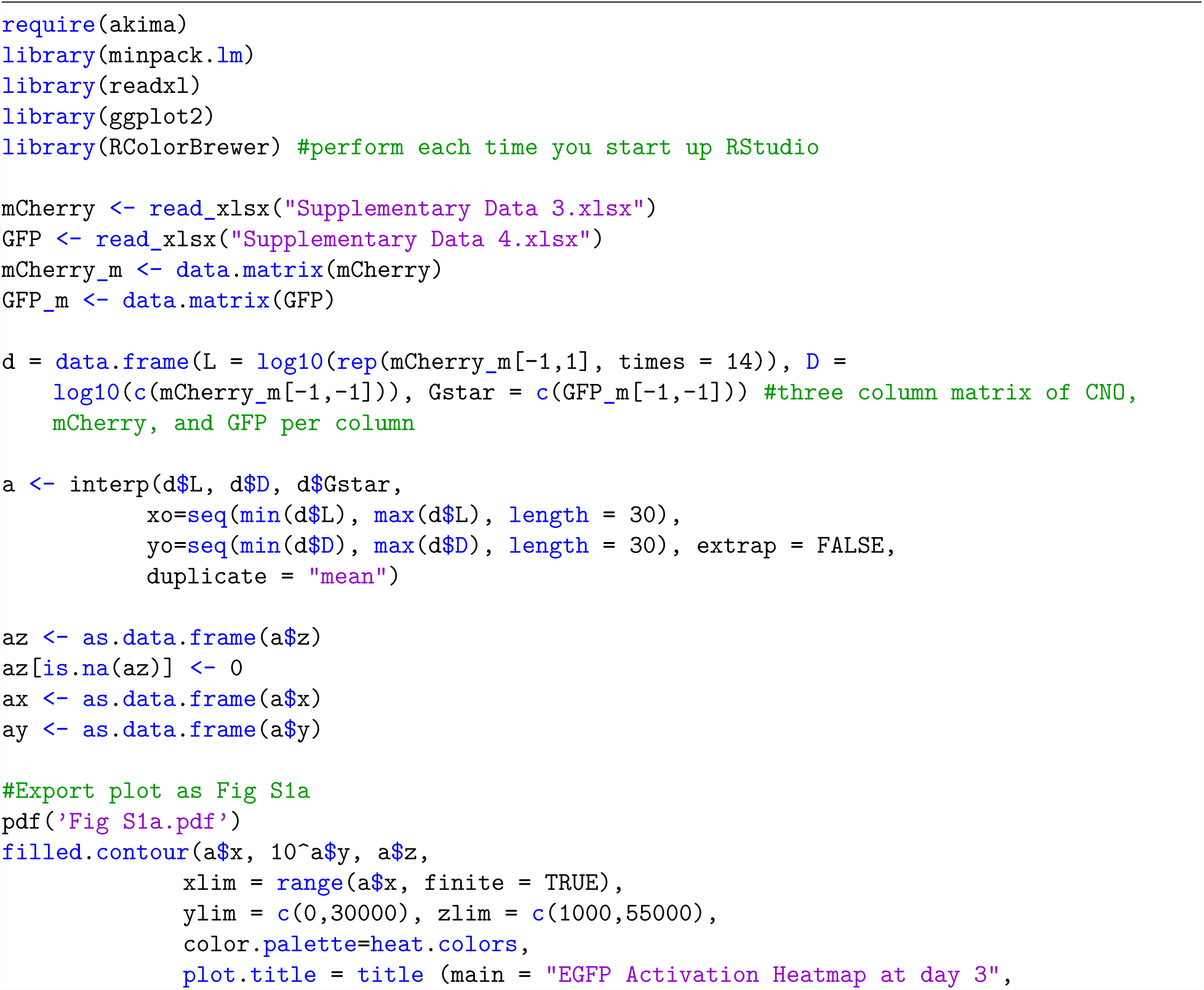
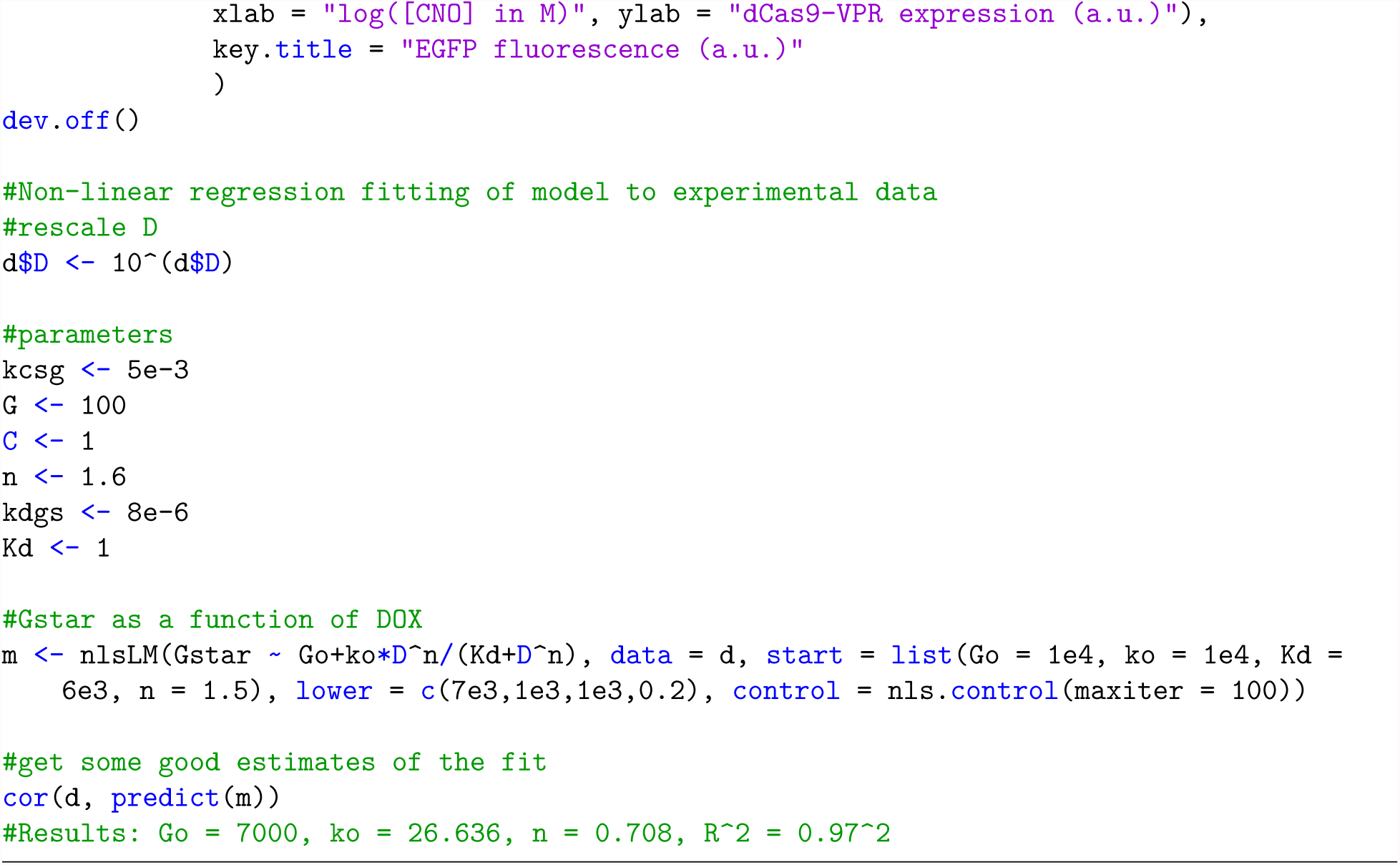

The purpose of the following R code is to plot Equation (6) in **Supplementary Note 2** with the fitted parameter values (**Fig. S1a, right**).

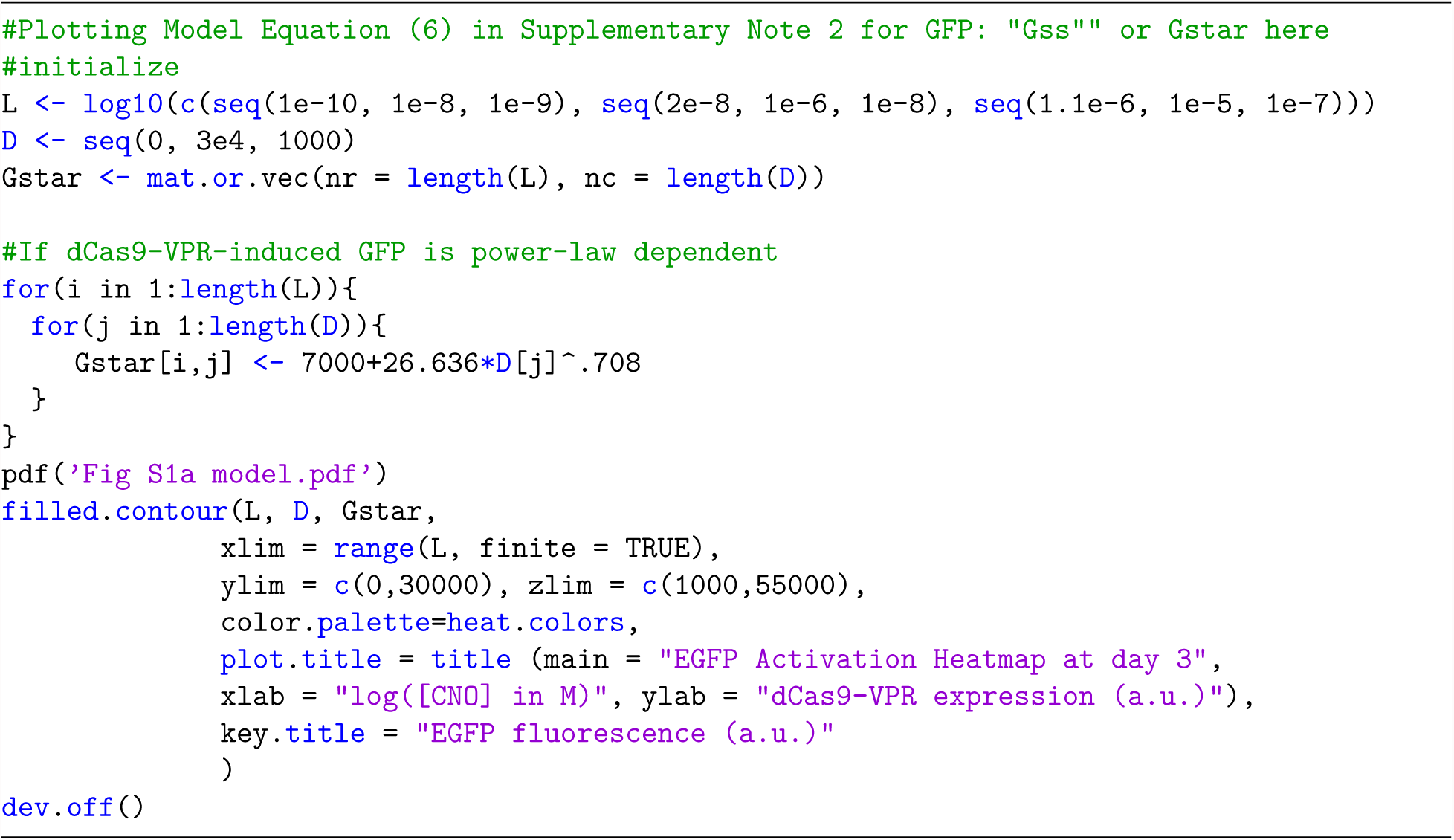

## Legends for Supplementary Movies

**Supplementary Movie 1 |** Representative time-lapse movie of hM3D-CRISPR ChaCha variant **K** in HEK293T TRE3G-EGFP reporter cells. Cells were imaged for 48 h, immediately after ±CNO stimulation (20μM) 1-day post-transfection. Upon CNO ligand treatment, hM3D-V2-TEVp-p2a-BFP (not shown) is expected to interact and cleave ARRB2-TCS-dCas9-VPR-mCherry. Released dCas9-VPR-mCherry then activates *EGFP*, translation of which was observed as early as 12 h after CNO treatment. Image frames were taken every 0.5 h. Movie rendered at 7 frames per second. PhC, Phase contrast. Scale bar, 50 μm.

**Supplementary Movie 2 |** Additional time-lapse fields of view of ARRB2-TCS-dCas9-VPRmCherry (note nuclear exclusion of mCherry) in HEK293T TRE3G-EGFP reporter cells. Cells were imaged for 48 h, immediately after ±CNO stimulation (20μM) 1-day post-transfection. Image frames were taken every 0.5 h. Movie rendered at 7 frames per second. Scale bar, 50 μm.

**Supplementary Movie 3 |** Additional time-lapse fields of view of EGFP reporter activation by hM3D-CRISPR ChaCha variant **K**, in HEK293T TRE3G-EGFP reporter cells. Cells were imaged for 48 h, immediately after ±CNO stimulation (20μM) 1-day post-transfection. Phase contrast. Image frames were taken every 0.5 h. Movie rendered at 7 frames per second. Scale bar, 50 μm.

## Legends for Supplementary Data

**Supplementary Data 1 |** ARRB2-dCas9-mCherry expression values, related to **Fig. 1c** and **Supplementary Fig. 1b**, and **Supplementary Note 2**.

**Supplementary Data 2 |** EGFP expression values, related to **Fig. 1c** and **Supplementary Fig. 1b**, and **Supplementary Note 2**.

**Supplementary Data 3 |** dCas9-mCherry expression values, related to **Fig. S1a** and **Supplementary Note 2**.

**Supplementary Data 4 |** EGFP expression values, related to **Fig. S1a** and **Supplementary Note 2**.

## REFERENCES

1. Lim, W.A., Designing customized cell signalling circuits. Nat Rev Mol Cell Biol, 2010. 11(6): p. 393–403.

2. Eshhar, Z., et al, Specific activation and targeting of cytotoxic lymphocytes through chimeric single chains consisting of antibody-binding domains and the gamma or zeta subunits of the immunoglobulin and T-cell receptors. Proc Natl Acad Sci U S A, 1993. 90(2): p. 720–4.

3. Morsut, L., et al, Engineering Customized Cell Sensing and Response Behaviors Using Synthetic Notch Receptors. Cell, 2016. 164(4): p. 780–791.

4. Lim, W.A. and C.H. June, The Principles of Engineering Immune Cells to Treat Cancer. Cell, 2017.168(4): p. 724–740.

5. Schwarz, K.A., et al, Rewiring human cellular input-output using modular extracellular sensors. Nat Chem Biol, 2017.13(2): p. 202–209.

6. Barnea, G., et al, The genetic design of signaling cascades to record receptor activation. Proceedings of the National Academy of Sciences of the United States of America, 2008. 105(1): p. 64–69.

7. Kroeze, W.K., et al, PRESTO-Tango as an open-source resource for interrogation of the druggable human GPCRome. Nat Struct Mol Biol, 2015. 22(5): p. 362–9.

8. Lauffenburger, D.A. and J.J. Linderman, Receptors: models for binding, trafficking, and signaling. 1996, New York: Oxford University Press. x, 365 p.

9. Katritch, V., V. Cherezov, and R.C. Stevens, Diversity and modularity of G protein-coupled receptor structures. Trends Pharmacol Sci, 2012. 33(1): p. 17–27.

10. Rosenbaum, D.M., S.G. Rasmussen, and B.K. Kobilka, The structure and function of G-protein-coupled receptors. Nature, 2009. 459(7245): p. 356–63.

11. Vardy, E., et al, A New DREADD Facilitates the Multiplexed Chemogenetic Interrogation of Behavior. Neuron, 2015. 86(4): p. 936–46.

12. Armbruster, B.N., et al, Evolving the lock to fit the key to create a family of G protein-coupled receptors potently activated by an inert ligand. Proc Natl Acad Sci U S A, 2007. 104(12): p. 5163–8.

13. Jinek, M., et al, RNA-programmed genome editing in human cells. Elife, 2013. 2: p. e00471.

14. Mali, P., et al, RNA-guided human genome engineering via Cas9. Science, 2013. 339(6121): p. 823–6.

15. Cong, L., et al, Multiplex genome engineering using CRISPR/Cas systems. Science, 2013. 339(6121): p. 819–23.

16. Gilbert, L.A., et al, CRISPR-mediated modular RNA-guided regulation of transcription in eukaryotes. Cell, 2013.154(2): p. 442–51.

17. Mali, P., et al, CAS9 transcriptional activators for target specificity screening and paired nickases for cooperative genome engineering. Nat Biotechnol, 2013. 31(9): p. 833–8.

18. Zalatan, J.G., et al, Engineering complex synthetic transcriptional programs with CRISPR RNA scaffolds. Cell, 2015.160(1-2): p. 339–50.

19. Konermann, S., et al, Genome-scale transcriptional activation by an engineered CRISPR-Cas9 complex. Nature, 2015. 517(7536): p. 583–8.

20. Hilton, I.B., et al, Epigenome editing by a CRISPR-Cas9-based acetyltransferase activates genes from promoters and enhancers. Nat Biotechnol, 2015. 33(5): p. 510–7.

21. Thakore, P. I., et al, Highly specific epigenome editing by CRISPR-Cas9 repressors for silencing of distal regulatory elements. Nat Methods, 2015. 12(12): p. 1143–9.

22. Barnea, G., et al, The genetic design of signaling cascades to record receptor activation. Proc Natl Acad Sci U S A, 2008.105(1): p. 64–69.

23. Chavez, A., et al, Highly efficient Cas9-mediated transcriptional programming. Nat Methods, 2015. 12(4): p. 326–8.

24. Sternberg, S.H., et al, DNA interrogation by the CRISPR RNA-guided endonuclease Cas9. Nature, 2014. 507(7490): p. 62–7.

25. Kapust, R.B., et al, The P1’ specificity of tobacco etch virus protease. Biochem Biophys Res Commun, 2002. 294(5): p. 949–55.

26. Sainz, E., et al, Four amino acid residues are critical for high affinity binding of neuromedin B to the neuromedin B receptor. J Biol Chem, 1998. 273(26): p. 15927–32.

27. Langenhan, T., et al, Model Organisms in G Protein-Coupled Receptor Research. Mol Pharmacol, 2015. 88(3): p. 596–603.

28. Wise, A., S.C. Jupe, and S. Rees, The identification of ligands at orphan G-protein coupled receptors. Annu Rev Pharmacol Toxicol, 2004. 44: p. 43–66.

29. Davila, M.L., et al, Efficacy and toxicity management of 19-28z CAR T cell therapy in B cell acute lymphoblastic leukemia. Sci Transl Med, 2014. 6(224): p. 224ra25.

## REFERENCES

1. Chavez, A., et al, Highly efficient Cas9-mediated transcriptional programming. Nat Methods, 2015. 12(4): p. 326–8.

2. Barnea, G., et al, The genetic design of signaling cascades to record receptor activation. Proceedings of the National Academy of Sciences of the United States of America, 2008. 105(1): p. 64–69.

3. Isberg, V., et al, GPCRdb: an information system for G protein-coupled receptors. Nucleic Acids Res, 2016. 44(D1): p. D356–64.

4. Horn, F., et al, GPCRDB: an information system for G protein-coupled receptors. Nucleic Acids Res, 1998. 26(1): p. 275–9.

5. Boc, A., A.B. Diallo, and V. Makarenkov, T-REX: a web server for inferring, validating and visualizing phylogenetic trees and networks. Nucleic Acids Res, 2012. 40(Web Server issue): p. W573–9.

